# Dual transcriptional analysis of *Peronospora belbahrii* and *Ocimum basilicum* in susceptible interactions

**DOI:** 10.1101/2021.09.07.457810

**Authors:** Eric T. Johnson, Hye-Seon Kim, Miaoying Tian, Nativ Dudai, Ofir Tal, Itay Gonda

## Abstract

Basil downy mildew, caused by the pathogen *Peronospora belbahrii*, is a major problem for sweet basil growers worldwide. The genome sequences of both *Ocimum basilicum* and *P. belbahrii* were recently completed but extensive transcriptome analysis of this pathosystem has not been completed. RNA sequencing was performed using basil leaf samples collected three and six days after inoculation with sporangia from an Illinois isolate of *P. belbahrii* and differentially expressed genes were identified. Gene enrichment analysis identified 22 genes that were upregulated at day three, in comparison to mock inoculated leaf samples, that were classified as ‘defense response to oomycetes’; among this group were five orthologues of *Arabidopsis thaliana DOWNY MILDEW RESISTANCE 6*. During the same time interval, many genes contributing to photosynthesis in the infected leaves were downregulated in comparison to control leaf samples. Many more genes were differentially expressed in the inoculated basil leaves at day six, compared to mock inoculated leaves, as the pathogen began to produce sporangiophores. On days three and six, the pathogen produced high transcript levels of secreted glycoside hydrolases, which likely release sugars from the plant cell walls needed for the growth of the pathogen. These results contribute to a better understanding of the infection process of downy mildew and will aid the development of more effective measures for reducing the severity of the disease.

## Introduction

Basil downy mildew, caused by *P. belbahrii*, is an obligate biotrophic oomycete. Outbreaks of this disease were first described in Switzerland in 2001 (Belbahri *et al.* 2005) and the disease was spread worldwide, likely due to contaminated seeds (Farahani-Kofoet *et al.* 2012). Annual losses due to this disease were estimated in the tens of millions of U.S. dollars (Wyenandt *et al.* 2015). Basil downy mildew was first found in the United States in 2007 (Roberts *et al.* 2009), and was detected in most states by 2014 (Wyenandt *et al.* 2015). Basil downy mildew can produce yellowing of the leaves and sporangia emerge from the stomata on the abaxial side of the leaves 7-10 days after the initial infection (Koroch *et al.* 2013; Farahani-Kofoet *et al.* 2014). The disease can be managed by fungicides (Homa *et al.* 2014; McGrath and Sexton 2017; Raid *et al.* 2017; Zhang *et al.* 2018). However, there are not effective control measures for growers of organic basil (Wyenandt *et al.* 2015). Alteration of growing conditions can limit the impact of downy mildew (Cohen *et al.* 2013; Cohen and Ben-Naim 2016). Genetic studies suggested that resistance to basil downy mildew can be acquired with 2 or more genes, but the functions of the resistance genes are not known (Pyne *et al.* 2017; BEN-NAIM *et al.* 2018).

The process by which *P. belbahrii* infects basil leaves is partially characterized at the anatomical level. *P. belbahrii* requires several hours of leaf wetness for infection (Garibaldi *et al.* 2007; Cohen and Ben-Naim 2016). The sporangia produce a single germ tube on the leaf surface that forms an appressorium which penetrates the epidermis (Zhang *et al.* 2019) but other laboratories have noted appressorium penetration in the stomatal opening (Koroch *et al.* 2013; Cohen *et al.* 2017). It is unknown if *P. belbahrii* germ tubes can absorb nutrients; if not, then growth would be limited to the energy reserves in the sporangium. Germ tubes of *Peronospora tabacina* grew 50 µm or more before penetration of tobacco callus cells (Trigiano *et al.* 1984) while the germ tubes were much shorter (10 µm) when penetrating tobacco leaves (Henderson 1937; McKeen and Svircev 1981). After penetration of the leaf, *P. belbahrii* hyphae grow intercellularly through mesophyll tissue; haustoria form on some mesophyll cells (Wyenandt *et al.* 2015). It is unknown if *P. belbahrii* hyphae can absorb nutrients, but haustoria are generally thought to be the nutrition acquisition structures of oomycetes (Hahn and Mendgen 2001). Regardless of which *P. belbahrii* anatomical structure absorbs nutrients, the carbohydrates and amino acids produced by basil must cross the plasma membrane of *P. belbahrii.* Carbohydrate acquisition in all organisms is partially facilitated by transmembrane proteins called sugar transporters (STs) and SWEETs (Sugars Will Eventually be Exported Transporters). ST proteins usually have 12 transmembrane helices while the SWEETs generally have 7 transmembrane helices (Abramson *et al.* 2003; Hu *et al.* 2016; Peng *et al.* 2018).

The genome of *P. belbahrii* was recently sequenced and annotated; a little over 9000 protein-coding genes were identified (Thines *et al.* 2020). The genomic data of *P. belbahrii* and other downy mildew pathogens indicate that their metabolic networks were generally smaller than oomycetes that are hemibiotrophs or necrotrophs (Spanu 2012; Rodenburg *et al.* 2018). For example, there were fewer metabolic reactions involved in sterol and phenylpropanoid biosynthesis in *P. belbahrii* than in *Phytophthora sojae, Phytophthora infestans* and *Phytophthora capsici* (Thines *et al.* 2020). In addition, *P. belbahrii* lacks enzymes for nitrate and nitrite assimilation, which means the pathogen likely acquires amino acids from the host to synthesize its own proteins (Thines *et al.* 2020). Bioinformatic analysis predicted 413 genes in *P. belbahrii* that encode for secreted proteins (Thines *et al.* 2020), which may be critical for pathogenesis. The genome of sweet basil ‘Perrie’, a Genovese-type cultivar, was recently sequenced and annotated (Gonda *et al.* 2020). The genome size of this tetraploid is 2.13 Gbp. With the genomes of the host and pathogen available, we performed a whole basil transcriptome analysis of detached basil leaves of a susceptible cultivar at three and six days after inoculation (dai) with *P*. *belbahrii* sporangia. The expression of the *P*. *belbahrii* genes coding for secreted proteins was completed at the same time points. The analysis identified several thousand differentially expressed basil genes in response to the pathogen while the genes of a number of secreted proteins were highly expressed by *P*. *belbahrii*.

## Materials and methods

### Conservation of *P*. *belbahrii* sporangia, Illinois

Seeds of Genovese basil (*Ocimum basilicum*, Johnny’s Selected Seeds, Winslow, Maine) were sown into seven punnets (13 x 13 cm) containing pasteurized soil (Sunshine Propagation Mix, Sungrow Horticulture, Canada) on a weekly basis. The punnets were placed into flat trays that were filled with water as needed and incubated in a growth chamber as previously described (Zhang *et al.* 2019). Liquid fertilizer (around 500 ml of Peter’s fertilizer at 1 g/l) was added to a plant tray five-seven days prior to harvest of leaves. Basil plants from Danville, Illinois with downy mildew symptoms were collected in 2014 and sporangia were sprayed onto young basil seedlings in the laboratory that were later used for inoculations of new seedlings as previously described (Zhang *et al.* 2019). A continuous supply of sporangia was produced through weekly inoculations of basil seedlings; a turbid suspension of sporangia (the concentration was not calculated) was sufficient to perpetuate the disease.

### Inoculation of detached leaves, Illinois

The first true basil leaves were excised from plants and treated with 10% bleach (0.825% sodium hypochlorite) for two minutes. The leaves were washed three times with sterile distilled water and placed onto sterile filter paper (with the adaxial side contacting the filter paper) inserted into Petri dishes (100 x 25 mm). *P*. *belbahrii* sporangia were collected into sterile water and the concentration diluted to 5 x 10^4^ sporangia ml^-1^. Approximately 300-500 µl of sporangial inoculation was added to the abaxial surface of each leaf. Control leaves received 300-500 µl of sterile water. The sterile filter paper was moistened with two ml of sterile water. All the dishes were kept in the dark at room temperature overnight; the covers on top of the dishes were sealed with parafilm. The following day the inoculum or sterile water was removed from each leaf and all the dishes (sealed with parafilm) placed in an incubator with 13 h of fluorescent lighting (11 h darkness) maintained at 25 °C and 33% relative humidity. Control and inoculated leaves were snap frozen in liquid nitrogen at three and six dai and stored at −80 °C.

### RNA extraction and transcriptome sequencing, Illinois

The RNA extractions were performed on individual leaves. For each treatment, at each time point, at least four leaves (from at least two plants) were extracted separately. There were five biological replicates of *P*. *belbahrii* inoculated leaves at three dai (labeled 3B) and five biological replicates of water inoculated leaves at three dai (labeled 3W). There were four biological replicates of *P*. *belbahrii* inoculated leaves at six dai (labeled 6B) and five biological replicates of water inoculated leaves at six dai (labeled 6W; details on all the samples are in Supplementary File 1). Frozen leaves were individually pulverized with a mortar and pestle using liquid nitrogen. Powdered tissue was transferred to a 1.5 ml centrifuge tube containing one ml of Trizol (Invitrogen, Carlsbad, CA). RNA was extracted using the Trizol Plus RNA Purification Kit (Invitrogen). Samples were treated with DNase (Qiagen, Hilden, Germany) and purified using the Gene Jet RNA Cleanup Kit (Thermo Fisher Scientific, Waltham, MA). RNA quality and quantity were assessed using a Nanodrop 2000 (Thermo Fisher Scientific) and a Tapestation 4200 (Agilent Technologies, Santa Clara, CA). All RNA samples had a RIN value > 6.0. The mRNA was purified from each RNA sample, converted to cDNA, and sequenced on an Illumina HiSeq or NovaSeq instrument (San Diego, CA) using a paired end 2 x150 bp read length (GENEWIZ, Plainfield, NJ). The final read numbers and mapping results (for all the transcriptomes) are available in Supplementary File 1. The raw read data of the Illinois experiments are deposited as PRJNA647030 in NCBI.

### Hawaii *P. belbahrii* isolate transcriptome experiments

A Hawaiian isolate of *P. belbahrii* was propagated on basil plants as previously described (Shao and Tian 2018). Four-week old basil Dolly plants were inoculated by spraying sporangial suspension (1 x 10^4^ sporangia ml^-1^) on the first pair of true leaves. Inoculated plants were kept in trays covered with plastic domes for maintaining high humidity and placed in the dark for 24 h. The plants were moved to a growth chamber with a 12 h photoperiod. The inoculated leaves were collected after three, four and five d, frozen in liquid nitrogen, and stored at −80 °C until total RNA extraction; a previous study indicated the Hawaiian *P. belbahrii* isolate significantly grew through Dolly basil leaves by three d (Shao and Tian 2018). RNA was isolated using RNeasy Plant Mini Kit (Qiagen) and contaminating gDNA was removed using DNA-free™ kit (Thermo Fisher Scientific). Libraries with 280 bp inserts were constructed using TruSeq RNA library Prep Kit v2 (Illumina), and the whole transcriptomes were sequenced using the 100 bp paired-end mode on a HiSeq 2500 (Illumina). The raw read files are deposited as PRJNA742033 in NCBI.

### Basil transcriptome analysis

Raw RNA-seq reads were filtered and cleaned using Trimmomatic (Bolger *et al.* 2014) to remove adapters and low-quality reads. Clean reads were mapped with STAR (Dobin *et al.* 2013) to the sweet basil genome (Gonda *et al.* 2020). Transcript quantification and differential gene expression analysis were performed using Cufflinks (v2.2.1) software (Roberts *et al.* 2011) combined with gene annotations. Genes were considered differentially expressed when the q-value was < 0.05 and expressed as log_2_ fold change (LFC); LFC values between −1 and 1 were discarded from the analysis. FPKM (Fragments Per Kilobase of transcript per Million fragments mapped) values of 35,011 genes (with expression value >1 in at least 4 transcriptome samples) were subjected to principal component analysis (PCA) with JMP software (SAS Institute Inc.) using the wide estimation method. The amino acid sequences of all the differentially expressed genes were analyzed by BLAST2GO using OmicsBox software. A Gene Matrix Transposed (GMT) file was built from a BLAST2GO output file in which all the basil differentially expressed genes associated with a specific GO term were listed; the GMT file was uploaded to g:Profiler (Raudvere *et al.* 2019). Ranked lists of upregulated and downregulated genes were submitted to g:Profiler for enrichment analysis of gene ontology (GO) terms (Raudvere *et al.* 2019).

### *P*. *belbahrii* transcriptome analysis

This analysis was completed using CLC Genomics Workbench version 12.0.3 (Qiagen, Valencia, CA) software. Raw sequencing reads (generated from Illinois and Hawaii transcriptomes, described above) were trimmed to remove adapter sequences and low quality reads (quality score limit 0.05 on the Phred scale). The “map reads to reference” function was used for mapping the reads to the *P*. *belbahrii* genome (GCA_902712285.1). The expression of the following putative *P*. *belbahrii* genes were examined: 21 sugar transporters; 22 housekeeping genes; and 411 secreted proteins (Thines *et al.* 2020). A gene expression value (TPM, transcripts per million) was calculated for each gene in each transcriptome sample. The data from the Illinois transcriptomes were normalized using the trimmed mean of M-values normalization method (Robinson and Oshlack 2010).

### Phylogenetic analysis and protein alignment

Proteins were aligned and phylogenetic trees constructed using MEGA 10 (Kumar *et al.* 2018). The percent identity matrix of proteins was aligned using Clustal 2.1 available on the EMBL-EBI website (Madeira *et al.* 2019).

## Results

### G I have and have enetic response of basil to *P*. *belbahrii*

Transcriptomes were generated from the detached basil leaves that were harvested three and six dai with the Illinois *P. belbahrii* isolate; control transcriptomes were generated from water (mock) inoculated detached leaves. *P. belbahrii* sporangia were visible on the detached leaves at six dai (Supplementary File 2). Transcriptome reads were mapped to the basil genome with the following percentages: 72-79% in 3W samples, 64-69% in 3B samples, 76-79% in 6W samples, and 49-60% in 6B samples (Supplementary File 1). A total of 11,890 basil genes were differentially expressed in the entire dataset (Figure 1A, Supplementary File 3). Only 194 basil genes were differentially expressed in all the treatments while the largest number of differentially expressed unique genes, 2742, was found in comparing 6W to 6B transcriptomes. Inoculation with *P. belbahrii* resulted in the expression of 707 basil genes at three dai (Figure 1A) that were not expressed in the other treatments. Conversely, 1055 basil genes were differentially expressed only in control leaves over three days (3W vs. 6W) indicating significant genetic changes to the leaves during this period. Further analysis found that slightly over half (51-59%) of the differentially regulated genes were upregulated in each of the four transcriptome comparisons (Figure 1B).

**Figure 1.**
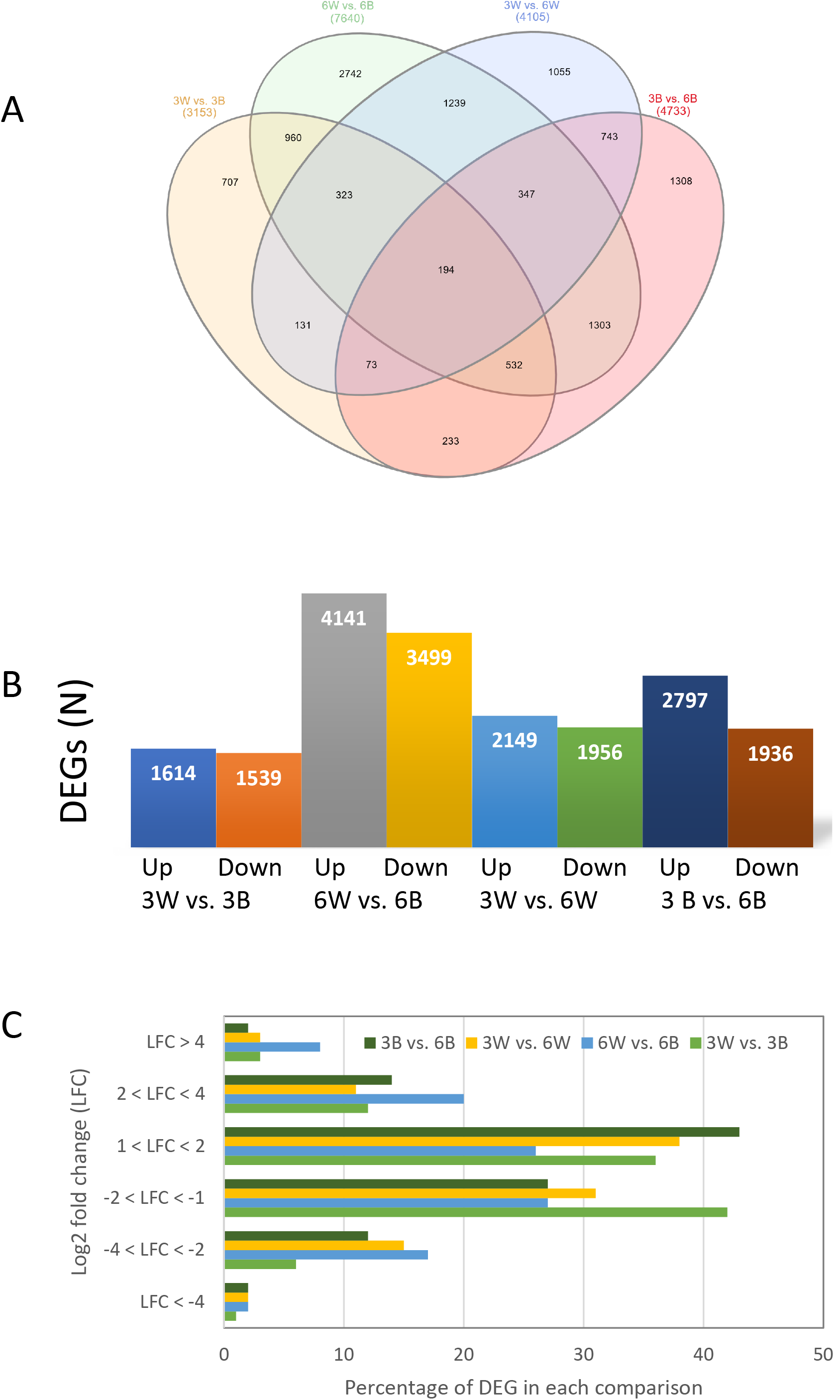
Venn diagram and expression levels of basil differentially expressed genes (DEGs). A. The Venn diagram shows the distribution of DEGs across the four comparisons. B. Each column shows the number of DEGs upregulated (up) or downregulated (down) with each comparison. C. The percentage of DEGs in each comparison grouped by six groups of expression levels.

The LFC values of the differentially expressed genes in each of the transcriptome comparisons were categorized into six different groups (Figure 1C). Examining all four comparisons, the expression levels of most of the genes were only modestly changed, with the majority of genes having LFC values between 1 and 2 or −1 and −2. For the 3W vs. 3B comparison, genes that had LFC values between −1 and −2 were the highest percentage (42%) of the LFC groups, but in the 3B vs. 6B comparison, genes that had LFC values between 1 and 2 were the highest percentage (43%) of the LFC groups. Interestingly, the largest number of differentially expressed genes with LFC values greater than 4 was found in the 6W vs. 6B comparison.

PCA was performed on the transcriptomes (Figure 2). Samples from the same treatment were generally clustered together. PC1 explained 31% of the variation in gene expression. Except for three samples, PC1 split all the samples into inoculated and non-inoculated samples. This result indicates that the overall gene expression was highly affected by the pathogen. When analyzed separately, a comparison between the PCA of 3B vs. 3W to the PCA of 6B vs. 6W (Supplementary File 4), suggests that the pathogen had a greater influence on gene expression at six dai. A comparison between PCAs of 3B vs. 6B and 3W vs. 6W (Supplementary File 4) suggests that the time of the sampling, evident in PC2, explains ∼21% of the variation, for both inoculated and non-inoculated samples.

**Figure 2.**
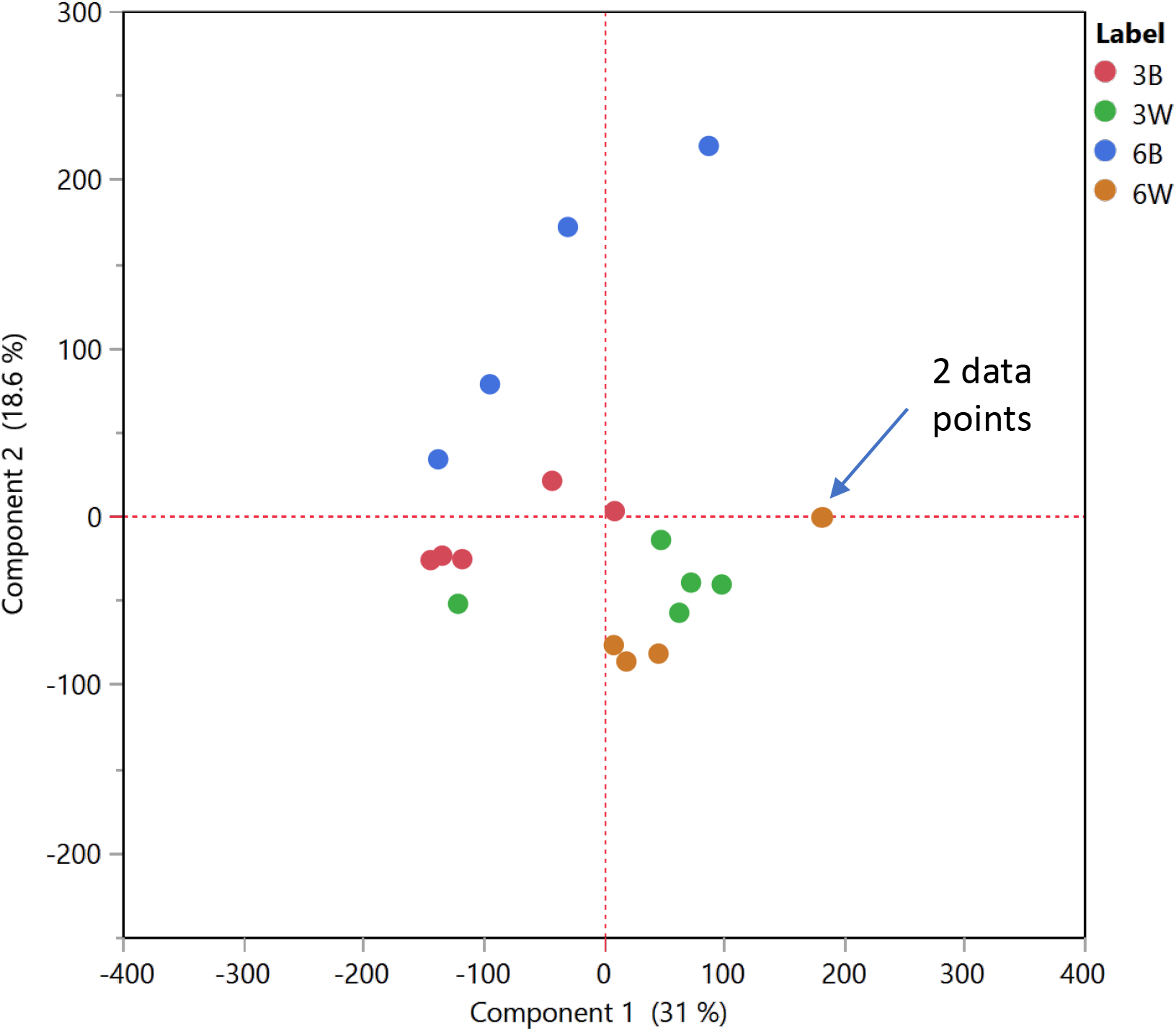
PCA analysis of basil FPKM values in all 19 transcriptomes.

### Basil genes upregulated in the 3B samples compared with 3W samples (3W vs. 3B)

Enrichment analysis utilizing Gene Ontology (GO) annotation was completed because of the large number of differentially expressed genes in the entire basil gene dataset. In the 3W v 3B comparison of upregulated genes there were 76 GO terms that were significant; ten of these terms are listed in Table 1 (also see Supplementary File 5). There were 22 genes that belonged to ‘defense response to oomycetes’ (listed in Supplementary File 5). The expression of these genes was induced in the range of 2.7-4.8 LFC. Five of these genes encode for naringenin, 2-oxoglutarate 3-dioxygenase (as annotated by BLAST2GO), an enzyme contributing to flavonoid biosynthesis, that belongs to the superfamily of the 2-oxoglutarate Fe(II) dependent oxygenases. Additional BLAST analysis of each of the five naringenin, 2-oxoglutarate 3-dioxygenase basil genes as queries with the *A. thaliana* genome determined that all were likely orthologues of At5g24530, the *DOWNY MILDEW RESISTANCE 6* (*DMR6*) gene, which is an 2-oxoglutarate Fe(II) dependent oxygenase that hydroxylates salicylic acid at the C5 position (Zhang *et al.* 2017). The five genes were 62-68% identical to the protein encoded by At5g24530 (Table 2). Analysis of the genomic locations of these genes found that two pairs of the five genes are located in homeologous scaffolds based on common BUSCO (Benchmarking Universal Single-Copy Orthologs) genes (Supplementary File 6) analyzed by (Gonda *et al.* 2020). However, XLOC_035244 possibly contains a point mutation, which results in a protein that is 79 amino acids shorter than XLOC_006768 at the C-terminus (Table 2). XLOC_006768 encodes a protein that is 339 amino acids, and the other three genes encode proteins that are all 336 amino acids. The 3W vs. 3B LFC of the five basil *DMR6-like* genes varied between 2.9-4.

**Table 1.** GO enrichment analysis in basil leaf gene expression.

| GO description | FDR | Number of genes |
| --- | --- | --- |
| **3W v 3B upregulated** |  |  |
| Integral component of endosome membrane | 3.4 E-12 | 18 |
| Calcium ion binding | 1.7 E-10 | 49 |
| Defense response to oomycetes | 1.2 E-9 | 22 |
| Regulation of nitric oxide metabolic process | 9.9 E-9 | 20 |
| Flavone biosynthetic process | 2.2 E-7 | 6 |
| Defense response to nematode | 2.2 E-6 | 14 |
| Cellular response to molecule of bacterial origin | 3.0 E-6 | 6 |
| Response to abiotic stimulus | 5.8 E-6 | 48 |
| L-aspartate transmembrane transporter activity | 8.0 E-6 | 10 |
| Cellular response to stress | 1.0 E-5 | 15 |
| **3W v 3B downregulated** |  |  |
| Chloroplast thylakoid membrane | 3.7 E-19 | 95 |
| Chloroplast stroma | 2.5 E-11 | 112 |
| mRNA binding | 6.6 E-9 | 109 |
| Oligosaccharide biosynthetic process | 2.9 E-6 | 7 |
| Galactinol-raffinose galactosyltransferase activity | 2.9 E-6 | 7 |
| Chloroplast nucleoid | 3.6 E-6 | 13 |
| Circadian rhythm | 4.3 E-6 | 51 |
| RNA processing | 9.1 E-6 | 19 |
| Raffinose catabolic process | 1.2 E-5 | 7 |
| Galactinol-sucrose galactosyltransferase activity | 3.6 E-5 | 7 |
| **6W v 6B upregulated** |  |  |
| Response to molecule of bacterial origin | 4.1 E-32 | 157 |
| Defense response to fungus | 3.1 E-24 | 230 |
| Response to chitin | 1.3 E-21 | 254 |
| Plasma membrane | 1.0 E-19 | 1065 |
| Protein autophosphorylation | 9.5 E-19 | 242 |
| Protein serine/threonine kinase activity | 5.1 E-18 | 213 |
| Response to salicylic acid | 2.1 E-17 | 149 |
| Defense response to oomycetes | 3.4 E-15 | 41 |
| Regulation of lignin biosynthetic process | 6.5 E-15 | 77 |
| Plant-type hypersensitive response | 8.0 E-11 | 62 |
| **6W v 6B downregulated** |  |  |
| Chloroplast thylakoid membrane | 9.0 E-32 | 229 |
| Chloroplast stroma | 2.2 E-29 | 300 |
| Chloroplast envelope | 7.2 E-27 | 265 |
| mRNA binding | 8.0 E-21 | 293 |
| Photosynthesis light harvesting in photosystem I | 3.8 E-19 | 34 |
| Chlorophyll binding | 1.2 E-16 | 46 |
| Plastoglobule | 1.5 E-16 | 71 |
| Apoplast | 1.5 E-14 | 218 |
| Photosystem I | 3.3 E-13 | 31 |
| Reductive pentose-phosphate cycle | 3.1 E-12 | 30 |

FDR - false discovery rate

**Table 2.** Properties of basil *DMR6*-like genes

| Gene | Length of protein | Homology to *A. thaliana* DMR6<sup>a</sup> | Differentially expressed in 3W vs. 3B<sup>b</sup> | Differentially expressed in 6W vs. 6B<sup>b</sup> |
| --- | --- | --- | --- | --- |
| XLOC_006768 | 339 | 62 | 3.9 | 6.2 |
| XLOC_024761 | 339 | 63 | NC | NC |
| XLOC_026367 | 336 | 67 | 2.9 | 4.1 |
| XLOC_035244 | 260 | 64 | 4.0 | 5.2 |
| XLOC_043494 | 336 | 68 | 3.0 | 5.1 |
| XLOC_058608 | 336 | 66 | 3.5 | 4.2 |
| XLOC_061828 | 336 | 68 | 2.6 | 3.8 |
| XLOC_071519 | 338 | 63 | NC | 5.8 |

<sup>a</sup> Comparison with protein sequences
<sup>b</sup> Values in column are LFC
NC is no change

XLOC_010371, one of the 22 genes in ‘defense response to oomycetes’ (Supplementary File 5), was annotated as a *DMR6-like oxygenase 1* gene by BLAST2GO; however, it is only 37-40% identical at the amino acid level to the five basil *DMR6-like* genes in ‘defense response to oomycetes’. BLAST analysis of XLOC_010371 using only the *A. thaliana* genome determined that it was closest to At2g36690, named *Germination Insensitive to ABA Mutant 2* (*GIM2*), which encodes a 2-oxoglutarate Fe(II) dependent oxygenase that has an essential role in seed germination in *A. thaliana* (Xiong et al. 2018). It is not clear how the basil orthologue of *GIM2* plays a role, if any, in plant defense to oomycetes; the 3W vs. 3B LFC of XLOC_010371 was 3.1.

Four of the genes of the ‘defense response to oomycetes’ term encode for calmodulin-binding protein 60 C (CBP60c); there are eight members of the CBP60 family in *A*. *thaliana*, some of which have roles in plant immunity (Truman *et al.* 2013). Other putative defense related genes in this group code for ABC transporter G 36 (ABCG36, one gene), WRKY transcription factor 70 (WRKY70, two genes), L-type lectin-domain containing receptor kinase (three genes), disease resistance protein RRS1 (one gene), a plant integral membrane protein (PIMP, two genes), and a transcription factor called SAR deficient 1 (involved in salicylic acid (SA) biosynthesis, two genes). This data indicate that the basil genome does contain a significant number of putative defense-related genes that are expressed in response to inoculation with *P. belbahrii*.

The GO terms ‘defense response to nematode’ and ‘cellular response to molecule of bacterial origin’ were also significantly enriched in the 3W vs. 3B upregulated comparison (Table 1). Six of the 14 genes in the term ‘defense response to nematode’ code for glucan endo-1,3-beta-D-glucosidase activity. 1,3-beta glucans contribute to the cell wall composition of *Phytophthora* and *Peronospora* oomycetes (MéLIDA *et al.* 2013), and therefore basil may use endo-1,3-beta-D-glucosidases as a weapon to defend against *P. belbahrii*. Genes encoding disease resistance protein RRS1 and WRKY70 were also in the ‘defense response to nematode’ term; different and identical genes for these same proteins were found in the ‘defense response to oomycetes’ term. Lastly, two genes encoding peroxidases were part of the ‘defense response to nematode’ term. There were six genes in the ‘cellular response to molecule of bacterial origin’, four of which encoded CBP60c and the other two encoded SAR deficient 1; all six genes were also part of the ‘defense response to oomycetes’.

In the 3W vs. 3B comparison of upregulated genes the term ‘integral component of endosome membrane’ had the lowest false discovery rate (FDR) value (Table 1). These proteins are partially embedded in the membrane, and all were capable of binding calcium. Not surprisingly, all 18 genes from the ‘integral component of endosome membrane’ term were also found in the ‘calcium ion binding’ term. It is possible that these proteins are involved in signaling pathways of basil defense from pathogens. The ‘regulation of nitric oxide metabolic process’ term contained 20 genes, all of which encoded proteins that were capable of binding calcium (as annotated by BLAST2GO). Eleven of the genes in ‘regulation of nitric oxide metabolic process’ were also found in the ‘integral component of endosome membrane’.

The ‘flavone biosynthetic process’ was another statistically significant GO term that contained six basil genes. Five of these genes were the same *DMR6-like* genes described above. The sixth gene (XLOC_061828), also a *DMR6* orthologous gene, was 99% identical to XLOC_043494 at the amino acid level; this pair are likely homeologous basil genes (Supplementary File 6). The 3W vs. 3B LFC of XLOC_061828 was 2.6.

There were a significant number of genes in the ‘response to abiotic stimulus’ term (Table 1). There were six E3 ubiquitin-protein ligase genes upregulated in this term; BLAST2GO annotation indicated the genes encode for four different proteins. The E3 ubiquitin-protein ligase gene family comprises at least 1000 members in *A. thaliana* some of which contribute to protein degradation and abiotic stress tolerance (Mazzucotelli *et al.* 2006). In addition, there were three 2-oxoglutarate (2OG)-Fe(II) oxygenase-like genes in this term. The ‘cellular response to stress’ term was significantly enriched in this analysis, possibly because of the change in abiotic conditions or due to the penetration of the downy mildew pathogen. Four of the 15 genes in this term coded for CBP60c.

### Basil genes downregulated in the 3B samples compared with 3W samples (3W vs. 3B)

In the 3W vs. 3B comparison of downregulated genes there were 65 GO terms that were significant; ten of these terms are listed in Table 1 (also see Supplementary File 5). The most significant terms were related to the chloroplast (thylakoid membrane and stroma) and ‘mRNA binding’, each term containing around 100 genes. The photosynthetic activity of the chloroplasts may have been reduced in order to shift metabolic resources to defense and stress tolerance. It is notable that ‘RNA processing’, related to ‘mRNA binding’, was also significantly enriched in this comparison. ‘Circadian rhythm’ gene expression, which is tied to photosynthetic gene expression, was also reduced three dai in the pathogen infected leaves. The terms ‘galactinol-raffinose galactosyltransferase activity’, ‘raffinose catabolic process’, ‘oligosaccharide biosynthetic process’ and ‘galactinol-sucrose galactosyltransferase activity’ were all significant in the 3W vs. 3B downregulated comparison. All four terms contained the same seven genes, which all encode for raffinose synthase. Raffinose family oligosaccharides are the primary carbohydrates transported in basil phloem, a symplastic phloem loader (Kang *et al.* 2007). The reduction in photosynthetic activity likely resulted in a parallel loss in raffinose synthesis.

### Basil genes upregulated in the 6B samples compared with 6W samples (6W vs. 6B)

In the 6W vs. 6B comparison of upregulated genes there were 253 GO terms that were significant (Supplementary File 5); ten of these terms are listed in Table 1. In general, there were more genes in the shared GO terms in the 6W vs. 6B upregulated comparison than the 3W vs. 3B upregulated comparison: ‘defense response to fungus’, 96 vs. 230 genes (3W vs. 3B upregulated comparison vs. 6W vs. 6B upregulated comparison); ‘response to chitin’, 63 vs. 254; ‘plasma membrane‘, 356 vs. 1065; ‘defense response to oomycetes’, 22 vs. 41; and ‘response to salicylic acid’, 59 vs. 149. This may reflect the fact that the gene expression in the pathogen infected leaves might be influenced by *P. belbahrii* to a greater degree at this time as the pathogen was likely growing throughout the entire leaf at six dai. In addition, ‘response to molecule of bacterial origin’, ‘defense response to fungus’ and ‘response to chitin’ were the three most significant terms in the 6W vs. 6B upregulated comparison. There were only 37 genes in common between ‘response to molecule of bacterial origin’ and ‘defense response to fungus’ and 50 genes in common between ‘defense response to fungus’ and ‘response to chitin’. Slightly over 1000 genes were part of the ‘plasma membrane’ term suggesting this cell compartment was the main stage of action between the host and the invaders. The terms ‘response to salicylic acid’ and ‘plant-type hypersensitive response’ were also highly significant; both are well known components of plant immunity.

The ‘defense response to oomycetes’ term in the 6W vs. 6B upregulated comparison had approximately twice the number of genes (41) as found in the 3W vs. 3B upregulated comparison (listed in Supplementary File 5); all 22 genes in the 3W vs. 3B upregulated comparison were also found in the 6W vs. 6B upregulated comparison. Some new genes added to the 6W vs. 6B upregulated comparison in the ‘defense response to oomycetes’ term included six genes encoding ABCG36; this gene is involved in nonhost resistance in *A. thaliana* (Stein *et al.* 2006). Seven basil genes that encode a DMR6-like protein were part of this group; five of the genes were the same ones identified in the ‘defense response to oomycetes’ term of the 3W vs. 3B upregulated comparison. Expression of one of the seven *DMR6-like* genes, XLOC_061828, was also induced in the 3W vs. 3B comparison that was found in the ‘flavone biosynthetic process’. Expression of the last *DMR6-like* gene, XLOC_071519, was not significantly induced in the 3W vs. 3B comparison; expression of its homeologous gene, XLOC_024761, was only induced in the 3B vs. 6B comparison at a statistically significant level. This study identified eight *DMR6-like* genes which can be grouped into 4 homeologous pairs (Supplementary File 6). Each pair is 95-98% identical at the amino acid level. The pairs XLOC_026367-XLOC_58606 and XLOC_043494-XLOC_061828 are fairly similar to each other (>86% amino acid identity among all 4 proteins). All eight of the basil DMR6-like amino acid sequences were included in a phylogenetic analysis of other DMR6 and DMR6-like oxygenases (such as those encoded by *DLO1* and *DLO2* from *A*. *thaliana*) from a few dicot plants (Figure 3 and Supplementary File 7). All eight basil DMR6-like proteins clustered in the clade with the *A*. *thaliana* DMR6 with a high percentage value (90) following a bootstrap test.

**Figure 3.**
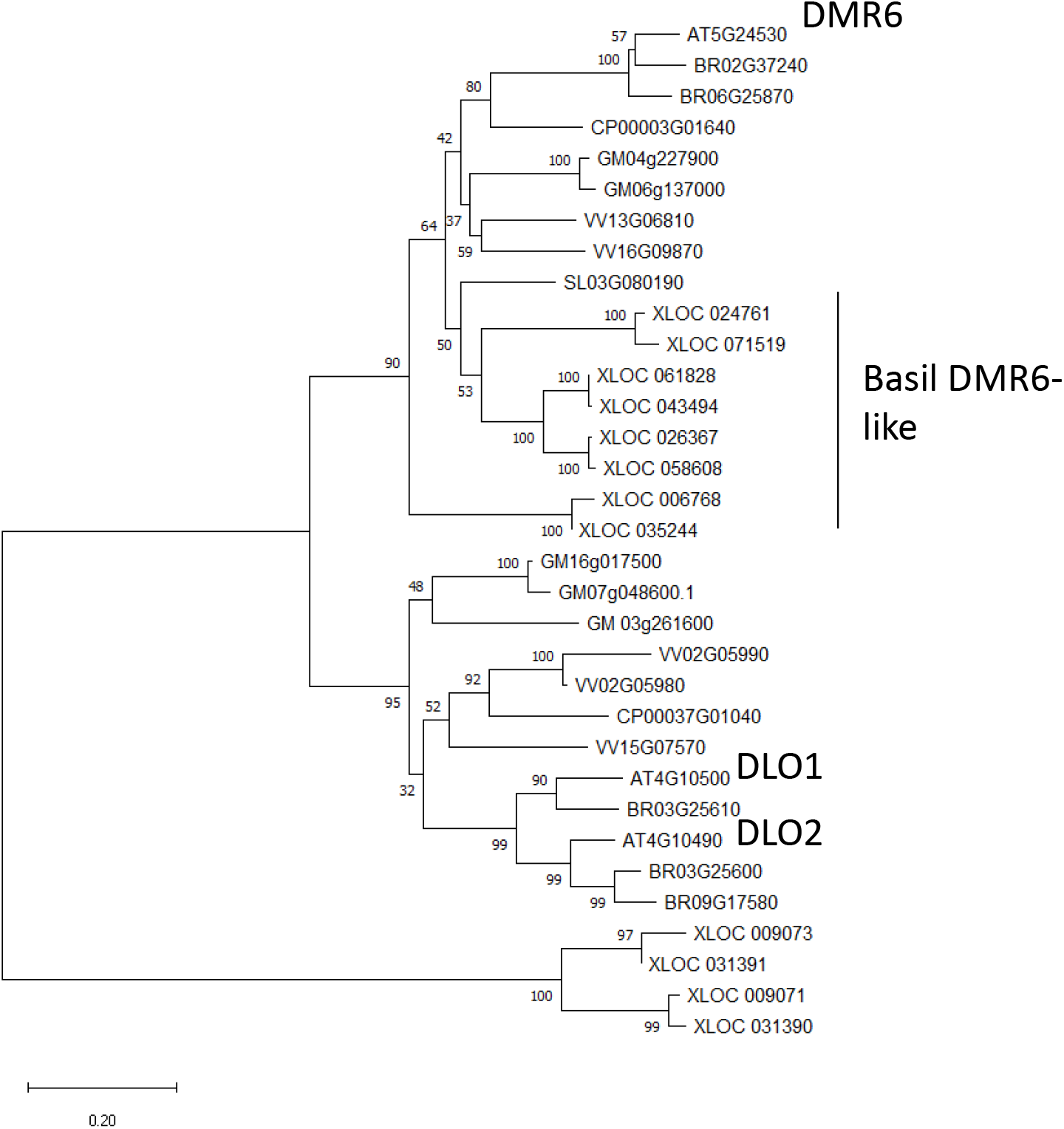
Phylogenetic tree of DMR6, DLO1 and DLO2 orthologous proteins in several dicot species based using the neighbor joining method. The percentage of replicate trees in which the associated taxa clustered together in the bootstrap test (1000 replicates) are shown next to the branches. The *A*. *thaliana* and *O. basilicum* DMR6-like proteins are noted. Other species included: BR, *Brassica rapa*; CP, *Carica papaya*; GM, *Glycine max*; SL, *Solanum lycopersicum*; VV, *Vitis vinifera*; XLOC, *Ocimum basilicum*

Another gene found in the 6W vs. 6B upregulated comparison in the ‘defense response to oomycetes’ term, XLOC_009073, was likely an orthologue of *A*. *thaliana DLO2* by BLAST analysis at the NCBI; XLOC_009073 was upregulated with a LFC of 3.8 (and upregulated with a LFC of 2.4 in the 3W vs. 3B comparison). The basil genome was queried for other genes related to XLOC_009073, and three others were found: XLOC_009071, XLOC_031390 and XLOC_031391. All these four basil genes encoded proteins were not related to other dicot genes encoding for DMR6 and DMR6-like oxygenases (Figure 3). In addition, only XLOC_009073 was differentially regulated in this study.

There were several of the 41 genes in the 6W vs. 6B upregulated comparison in the ‘defense response to oomycetes’ term (not upregulated at 3 dai) that might have contributed to defense against oomycetes based on research in other pathosystems: an additional WRKY70 (XLOC_051131); L-type lectin-domain containing receptor kinase (XLOC_054644); receptor like protein 7 (XLOC_058288); Cf-9 receptor (XLOC_0 49189); EIX2 receptor (XLOC_063298 and XLOC_063300); and G-type lectin S-receptor-like threonine protein kinase (XLOC_009955). Additional studies are needed to ascertain the role of these basil genes in plant defense.

### Basil genes downregulated in the 6B samples compared with 6W samples (6W vs. 6B)

In the 6W vs. 6B comparison of downregulated genes there were 132 GO terms that were significant; ten of these terms are listed in Table 1 (also see Supplementary File 5). Most of the highly significant terms in this comparison were involved with photosynthesis, except for the ‘mRNA binding’ and ‘apoplast’ terms. There were more downregulated genes in the ‘mRNA binding’ term in the 6W vs. 6B comparison than in the downregulated genes in the 3W vs. 3B comparison (293 vs. 109 genes). This might be due to the reduction in the gene expression devoted to photosynthesis. In addition, 23 of the GO terms in the downregulated genes of the 3W vs. 3B comparison were also part of the GO terms in the 6W vs. 6B comparison; 14 of the 23 terms involve photosynthesis or carbohydrate production. Slightly over 200 genes were downregulated in the apoplast, the anatomical compartment where much of the downy mildew growth has likely occurred at this time after inoculation.

The GO enrichment analysis favors genes with high LFC values. However, there may be basil genes with a moderate change in expression level that contribute to the susceptibility of Genovese basil to *P. belbahrii* but were not identified. On the other hand, there may be basil resistance genes that have been missed in this analysis. In addition, these experiments only examined changes in mRNA levels; epigenetic and posttranscriptional mechanisms of regulation are likely contributing to the defense response (which ultimately fails) of Genovese basil to *P. belbahrii*.

### *P. belbahrii* gene expression in basil

The leaf transcriptome samples inoculated with *P. belbahrii* were mapped to the *P. belbahrii* genome: 14-19% of reads in the Illinois 3B samples and 25-40% in the Illinois 6B samples (Supplementary File 1). In addition, three basil leaf transcriptomes were produced three, four and five d after inoculation with sporangia from a *P. belbahrii* isolate from Hawaii; mapping of these transcriptomes found that 11, 32 and 40% of the transcripts, respectively, were from *P. belbahrii* (Supplementary File 1).

An analysis of the expression of *P. belbahrii* genes encoding 411 secreted proteins (Thines et al. 2020) was completed (Supplementary Files 8 and 9). Some of the most highly expressed transcripts from this group coded for glycoside hydrolases (GH), though there were several highly-expressed transcripts that coded for proteins with no known function. All of the expression levels of the transcripts encoding secreted cell wall degrading enzymes were examined; those with mean transcripts per 1,000,000 reads (TPM) levels of 1000 or more are listed in Table 3 (genes with TPM values greater than 1,000 in all three Hawaii transcriptomes were also listed). The choice of 1000 TPM is based on the expression levels of several “housekeeping” genes (see below). Two *P. belbahrii* glycoside hydrolase genes with high expression levels in the Illinois experiment at three and six dai (PEBEL_30158 and PEBEL_30235) were highly expressed in the Hawaii experiment. Several *P. belbahrii* glycoside hydrolase genes (PEBEL_00769, PEBEL_03107.2, PEBEL_03490, PEBEL_04261) were well expressed in the Hawaii leaf transcriptomes but not in the Illinois leaf transcriptomes. These variations in gene expression may reflect differences between the Hawaii and Illinois experimental conditions or be due to genetic differences between the Hawaii and Illinois *P. belbahrii* pathogens. The PEBEL_04261 encoded protein contained motifs for GH17 as well as the UL36 domain, which will be described in more detail below.

**Table 3.** Expression of *P. belbahrii* genes in transcriptomes that encode secreted glycoside hydrolase proteins

| Gene | Illinois 3B N = 5 | Illinois 6B N = 4 | Hawaii leaves 3-5 d<br>N = 3 |
| --- | --- | --- | --- |
| PEBEL_00052 GH1 | 1637 ± 121 | 3419 ± 319 | 1743, 1974, 2657 |
| PEBEL_00769 GH3 | 2308 ± 281 | 735 ± 170 | 13554, 12425, 4929 |
| PEBEL_03107.2 GH37 | 378 ± 26 | 445 ± 89 | 1479, 1834, 2208 |
| PEBEL_03490 GH5 | 1601 ± 160 | 1763 ± 337 | 4989, 15831, 25834 |
| PEBEL_04261 GH17 +<br>UL36 domain | 1144 ± 221 | 2018 ± 735 | 20987, 21281,<br>13080 |
| PEBEL_04311 GH17 | 74417 ± 5412 | 100312 ± 17226 | 14093, 11238, 7870 |
| PEBEL_05385 GH 32 | 3472 ± 513 | 1121 ± 286 | 9975, 9185, 8082 |
| PEBEL_08766 GH6 | 23 ± 7 | 1480 ± 167 | 10, 18, 64 |
| PEBEL_30158 GH12 | 139826 ± 22669 | 132725 ± 22936 | 58392, 60913,<br>75906 |
| PEBEL_30235 GH12 | 98961 ± 5409 | 76210 ± 4514 | 34035, 46766,<br>30629 |

All values are TPM ± SE; GH is glycoside hydrolase

Gene expression was also examined for a number of *P. belbahrii* genes encoding for secreted virulence proteins, as previously annotated (Thines *et al.* 2020). Several genes that coded for putative effectors were expressed at moderate levels in both the Illinois and Hawaii transcriptomes (Table 4); six of the seven effector genes encoded for proteins containing a RxLR or RxLK motif, while PEBEL_07289 also contained a Nudix hydrolase domain (Dong and Wang 2016). RxLR effectors have been characterized as virulence factors in a number of different oomycetes (Deb *et al.* 2018; Liu *et al.* 2018; Ai *et al.* 2020). Proteins with a Nudix hydrolase domain have been identified in a number of plant pathogens, but their role is not well understood (Dong and Wang 2016). The PEBEL_01603 encoded effector contains both the DUF2403 and DUF2401 domains, which possibly enables the hydrolysis of β (Steczkiewicz *et al.* 2010). The gene PEBEL_00246.2 was well expressed in the Illinois transcriptomes at three dai, but expression declined after this time (Table 4). The protein encoded by PEBEL_00246.2 contains a necrosis inducing *Phytophthora* protein (NPP1) domain; proteins with this domain are also called NLPs (necrosis and ethylene inducing peptide 1-like proteins). NLPs occur in a variety of fungi, bacteria and oomycetes but the mechanism of the necrotic activity in plants caused by NLPs is not clearly known (Oome and Van den Ackerveken 2014).

**Table 4.**
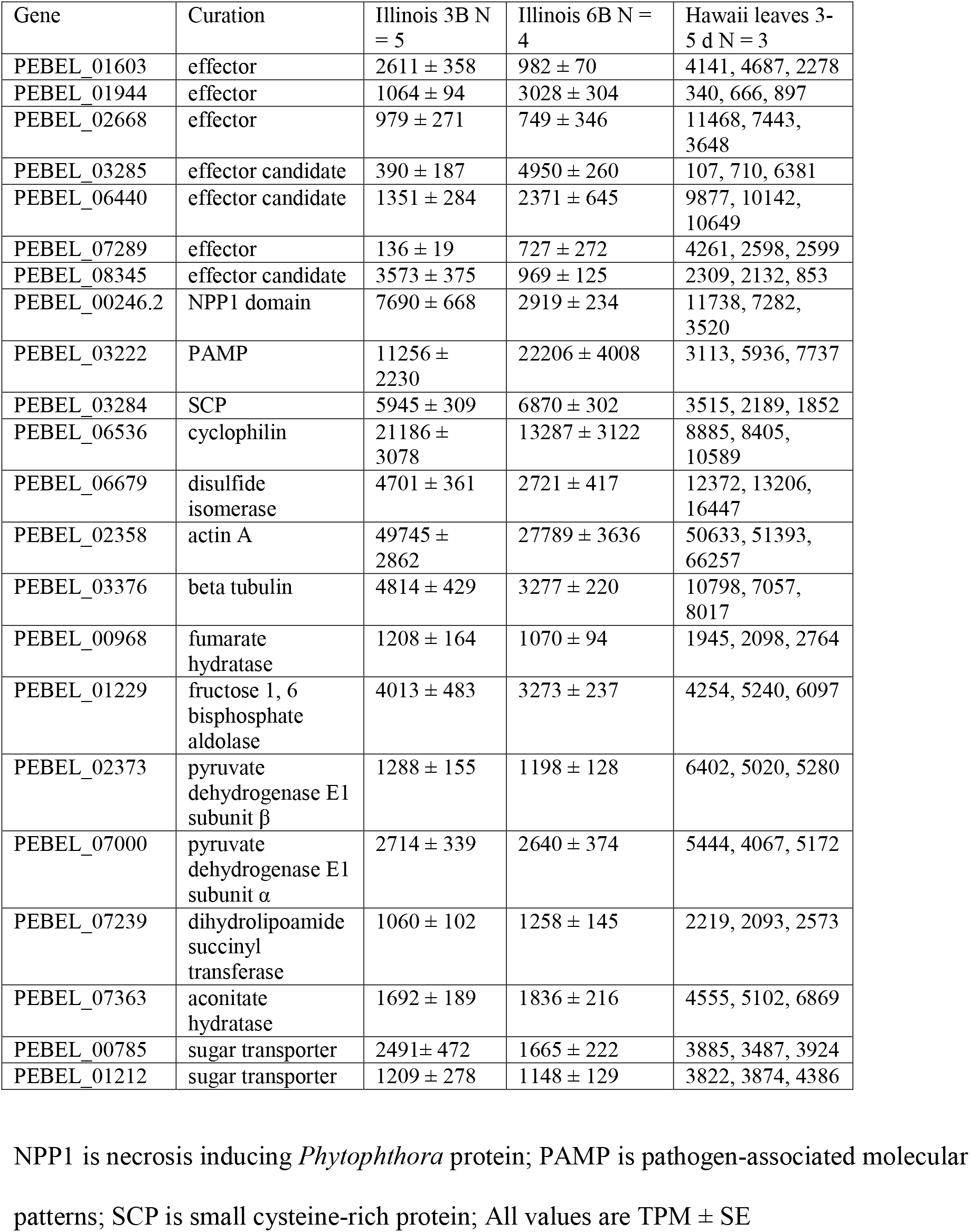
Expression of *P. belbahrii* genes in transcriptomes that encode putative virulence, structural and metabolic proteins

| Gene | Curation | Illinois 3B N = 5 | Illinois 6B N = 4 | Hawaii leaves 3-5 d N = 3 |
| --- | --- | --- | --- | --- |
| PEBEL_01603 | effector | 2611 ± 358 | 982 ± 70 | 4141, 4687, 2278 |
| PEBEL_01944 | effector | 1064 ± 94 | 3028 ± 304 | 340, 666, 897 |
| PEBEL_02668 | effector | 979 ± 271 | 749 ± 346 | 11468, 7443, 3648 |
| PEBEL_03285 | effector candidate | 390 ± 187 | 4950 ± 260 | 107, 710, 6381 |
| PEBEL_06440 | effector candidate | 1351 ± 284 | 2371 ± 645 | 9877, 10142, 10649 |
| PEBEL_07289 | effector | 136 ± 19 | 727 ± 272 | 4261, 2598, 2599 |
| PEBEL_08345 | effector candidate | 3573 ± 375 | 969 ± 125 | 2309, 2132, 853 |
| PEBEL_00246.2 | NPP1 domain | 7690 ± 668 | 2919 ± 234 | 11738, 7282, 3520 |
| PEBEL_03222 | PAMP | 11256 ± 2230 | 22206 ± 4008 | 3113, 5936, 7737 |
| PEBEL_03284 | SCP | 5945 ± 309 | 6870 ± 302 | 3515, 2189, 1852 |
| PEBEL_06536 | cyclophilin | 21186 ± 3078 | 13287 ± 3122 | 8885, 8405, 10589 |
| PEBEL_06679 | disulfide isomerase | 4701 ± 361 | 2721 ± 417 | 12372, 13206, 16447 |
| PEBEL_02358 | actin A | 49745 ± 2862 | 27789 ± 3636 | 50633, 51393, 66257 |
| PEBEL_03376 | beta tubulin | 4814 ± 429 | 3277 ± 220 | 10798, 7057, 8017 |
| PEBEL_00968 | fumarate hydratase | 1208 ± 164 | 1070 ± 94 | 1945, 2098, 2764 |
| PEBEL_01229 | fructose 1, 6 biphosphate aldolase | 4013 ± 483 | 3273 ± 237 | 4254, 5240, 6097 |
| PEBEL_02373 | pyruvate dehydrogenase E1 subunit β | 1288 ± 155 | 1198 ± 128 | 6402, 5020, 5280 |
| PEBEL_07000 | pyruvate dehydrogenase E1 subunit α | 2714 ± 339 | 2640 ± 374 | 5444, 4067, 5172 |
| PEBEL_07239 | dihydrolipoamide succinyl transferase | 1060 ± 102 | 1258 ± 145 | 2219, 2093, 2573 |
| PEBEL_07363 | aconitate hydratase | 1692 ± 189 | 1836 ± 216 | 4555, 5102, 6869 |
| PEBEL_00785 | sugar transporter | 2491 ± 472 | 1665 ± 222 | 3885, 3487, 3924 |
| PEBEL_01212 | sugar transporter | 1209 ± 278 | 1148 ± 129 | 3822, 3874, 4386 |

NPP1 is necrosis inducing *Phytophthora* protein; PAMP is pathogen-associated molecular patterns; SCP is small cysteine-rich protein; All values are TPM ± SE

The PEBEL_03222 elicitin like gene (also called a pathogen-associated molecular pattern, PAMP) was highly expressed in the Illinois *P. belbahrii* transcriptomes, but not in the Hawaii *P. belbahrii* transcriptomes (Table 4). The PEBEL_03222 transcript encodes a protein with similarity to the elicitin-like protein SOL11B of *Phytophthora sojae* (E value was 9e^-46^ using BLASTP on NCBI); SOL11B belongs to the ELL-11 clade, whose members have a different cysteine spacing pattern than the typical elicitin (Jiang *et al.* 2006). The PEBEL_03284 transcript was moderately expressed in the Illinois and Hawaii transcriptomes. PEBEL_03284 encodes for small cysteine-rich protein (SCP); SCPs from several *Phytophthora* have been partially characterized as phytotoxic molecules (Orsomando *et al.* 2011; Zhang *et al.* 2021). Cyclophilin protein encoded by PEBEL_06536 was highly expressed in the Illinois transcriptomes, and moderately expressed in the Hawaii transcriptomes. Cyclophilins have demonstrated roles as pathogenicity factors in true fungi, but their function in oomycetes is not yet clear (Singh *et al.* 2018). The gene for disulfide isomerase (PEBEL_06679) was well expressed in the Hawaii transcriptomes but downregulated from three to six dai in the Illinois transcriptomes. The PEBEL_06679 encoded protein is very similar to the *Phytophthora parasitica* disulfide isomerase, which was shown to be a virulence factor infection in *Nicotiana benthamiana* (Meng *et al.* 2015).

The expression of *P. belbahrii* genes likely contributing to metabolism and structure of the pathogen were also examined (Table 4). The actin gene was highly expressed in all the transcriptomes but the number of beta tubulin transcripts were substantially less; there was a statistically significant reduction in the amount of beta tubulin transcripts when comparing the values of 3B vs. 6B, but the log 2 value (−1.0) is probably not biologically relevant (Gonçalves *et al.* 2019). Other genes contributing to metabolism were generally in the range of 1000-7000 TPM. We identified 21 genes in *P. belbahrii* likely to encode transmembrane transporters of carbohydrates (Supplementary File 11). Most of the carbohydrate transporter gene expression levels were lower than that of the genes contributing to metabolism listed in Table 3. The three carbohydrate transporter genes with the highest expression levels in the Illinois transcriptomes are listed in Table 3. One gene encodes for a SWEET transporter (Hu *et al.* 2016). The other two carbohydrate transporter genes encode for putative major facilitator superfamily transporters containing 12 predicted transmembrane helices (Chen *et al.* 2015).

## Discussion

### Genetic response of basil to *P*. *belbahrii*

The expression of a number of defense-related genes basil were induced by *P*. *belbahrii* three dai. At the same time, the number of transcriptome reads in leaves infected by the pathogen were 9-23% of the total transcriptome reads, indicating that the pathogen had substantially grown through the leaf, which was expected in this compatible pathogen-host interaction. It is possible that many of the resistance-related genes found in this study of a compatible interaction would also be expressed in an incompatible interaction.

L-type lectin-domain containing receptor kinases are found in the plant plasma membrane and possibly mediate adhesions to the cell wall; a mutation in one L-type lectin-domain containing receptor kinase led to increased susceptibility in *A*. *thaliana* to *Phytophthora brassicae* (Bouwmeester *et al.* 2011). Fourteen L-type lectin-domain containing receptor kinases in *A*. *thaliana* contribute to resistance against various *Phytophthora* (Wang and Bouwmeester 2017). L-type lectin-domain containing receptor kinases possibly contribute to oomycete resistance in a number of crop plants (Wang *et al.* 2015; Woo *et al.* 2016).

In *A*. *thaliana*, WRKY70 is a positive regulator of resistance to *Hyaloperonospora parasitica* and *Erysiphe chicoracearum* (LI et al. 2006; KNOTH et al. 2007). WRKY70 possibly activates the expression of genes involved in plant defense in *A*. *thaliana*, including *PR1, PR2, PR5* and *SARD1* (Li *et al.* 2004). Recent work determined that WRKY70 is phosphorylated after infection with *Pseudomonas syringae* pv. tomato DC3000 in *A*. *thaliana*, which activates the expression of *SARD1* (Liu et al. 2021). *SARD1* positively regulates SA biosynthesis in *A*. *thaliana*, which is a well-known plant defense signaling pathway (Wang *et al.* 2011). Further studies will need to be completed to see if these molecular signaling pathways working in a bacteria pathogen-*A*. *thaliana* system also operate in the basil-downy mildew interaction.

CBP60a is a repressor of plant immunity while CBP60g enhances plant immunity (Truman *et al.* 2013). In the same study, the CBP60c mutant of *A*. *thaliana* enhanced the growth of *Pseudomonas syringae* strain ES4326 (Truman *et al.* 2013). CBP60c may have a role in basil plant immunity which needs further characterization.

The loss of *PENETRATION3*, also called ABCG36, can change plant susceptibility to nonadapted and adapted pathogens, reduce the hypersensitive response and diminish race-specific disease resistance (Kobae *et al.* 2006; Stein *et al.* 2006; Loehrer *et al.* 2008; Xin *et al.* 2013; Johansson *et al.* 2014). PENETRATION3-GFP fusion protein was found at the plasma membrane in uninfected cells but was concentrated at pathogen infection sites in diseased leaves (Stein *et al.* 2006). Basil might be synthesizing ABCG36 for accumulation at sites of downy mildew infection. PENETRATION3 might be contributing to the transfer of toxic molecules at pathogen invasion sites (Stein *et al.* 2006).

The expression of *CaPIMP1*, which encodes a putative integral membrane protein, was induced by compatible and incompatible strains of *Xanthomonas campestris* pv. *vesicatoria* on pepper leaves (Hong *et al.* 2008). In *A*. *thaliana* plants overexpressing *CaPIMP1*, disease resistance to *Pseudomonas syringae* pv. tomato DC3000 was enhanced compared to wild type plants; in contrast, the *CaPIMP1* overexpressing plants were more susceptible to *H*. *arabidopsidis*, a biotrophic oomycete (Hong *et al.* 2008). The expression of the gene for another disease resistance protein, RRS1, was induced at both three and six dai. RRS1 pairs with RPS4 to form an immune complex; the complex contributes to resistance of bacterial and fungal invaders, but there is no published data regarding the complex and oomycete resistance (Huh *et al.* 2017; Castel *et al.* 2019). More experiments need to be completed to determine the contributions of RRS1 and PIMP1 to basil plant immunity.

This transcriptomic study indicates that basil contains a gene family of *DMR6-like* genes with at least eight members. The expression of six of these genes were well induced in basil leaves at both three and six dai. Six *DMR6-like* genes were identified in the genome of basil cultivar Genoveser; the authors indicate there were likely additional *DMR6-like* genes in the genome that they could not identify (Hasley *et al.* 2021). The expression of the single *DMR6* gene in *A*. *thaliana* was induced by both compatible and incompatible *H*. *parasitica* isolates in leaves; *DMR6* transcript levels increased 25-fold in the incompatible interaction at three dai, but the level of *DMR6* expression was induced almost 40-fold in the compatible reaction at four dai (Van Damme *et al.* 2008). Mutation of *DMR6* results in plants that are more resistant to *H*. *arabidopsidis* (Van Damme et al. 2008), that is likely due to elevated SA levels (Zhang *et al.* 2017), which enhances plant defense gene expression. No studies to date have examined the possibility that *P. belbahrii* manipulates the induction of a basil *DMR6-like* gene to enhance infection. Each basil DMR6-like protein may have to catalyze a unique substrate. Two *DMR6* genes in potato were individually mutated using CRISPR-Cas9, but only the mutation in *StDMR6-1* resulted in plants that had increased resistance against late blight (Kieu *et al.* 2021). This suggests that similar *DMR6* genes may have different functions. Identifying the function of each *DMR6-like* gene in basil could be helpful in developing a basil mutant that is mutated at the *DMR6-like* gene that hydroxylates SA, which might be resistant to downy mildew. Mutations at *DMR6*, or RNAi of *DMR6* expression, was successful in making disease resistant tomato and potato (de Toledo Thomazella *et al.* 2016; Sun *et al.* 2016; Kieu *et al.* 2021). On the other hand, mutation of a conserved region of all known basil *DMR6-like* genes using CRISPR-Cas9 resulted in basil plants that exhibited a reduction of *P. belbahrii* biomass and numbers of sporangia produced after inoculation of leaves with the pathogen, compared with wild type plants (Hasley *et al.* 2021). These studies indicate that *DMR6* is a good target for promoting disease resistance in plants.

### *P. belbahrii* gene expression in basil leaves

Three days after inoculation, the downy mildew pathogen was synthesizing a significant number of transcripts that are likely processed into cell wall degrading enzymes. These enzymes probably enable the pathogen to invade plant tissue and also provide nutrients for continued growth throughout the leaf (Walton 1994; Hématy *et al.* 2009). The possible substrates for GHs encoded in the *Phytophthora parasitica* genome were predicted by Blackman et al. It is likely that some GH enzymes have multiple substrates. The predicted substrates for the GHs of *P. belbahrii* (based on the data of Blackman et al. 2014) and their relative mRNA abundance in basil leaves (based on the transcriptome data in this study) are depicted in Figure 4. Cellulose content was 15-16% (w/w) of the dry biomass of cultured basil cells (Dalton 1984); cellulose can be digested by five different GHs (Blackman *et al.* 2014). In the basil-downy mildew susceptible interaction, most of the cellulose and hemicellulose is likely digested by the two enzymes with GH12 activity. Based on mRNA expression levels and substrate specificity, pectin digestion is likely minimal during the susceptible interaction. The GH17 substrate is β-1,3 glucan, the primary component of callose, which is sometimes synthesized by plants at pathogen infection sites (Hückelhoven 2007; Chen and Kim 2009). It seems that most of the mRNA for GH17 enzyme activity in the Illinois interaction was contributed only by PEBEL_04311 while it was split between PEBEL_04311 and PEBEL_04261 in the Hawaii basil-downy mildew interaction. In addition, the β-1,3 glucan could be digested by GH5 enzyme; PEBEL_03490 mRNA levels were significantly expressed in the Hawaii *P. belbahrii* transcriptomes. Lastly, sucrose and trehalose are also possible substrates in the downy mildew-basil susceptible interaction, but the predicted enzyme activity was not likely significant in the Hawaii and Illinois transcriptomes. These data suggest that *P. belbahrii* primarily digests cellulose and hemicellulose to obtain sugars from basil; the pathogen possibly prevents the plant from depositing callose, which will enhance pathogen growth.

**Figure 4.**
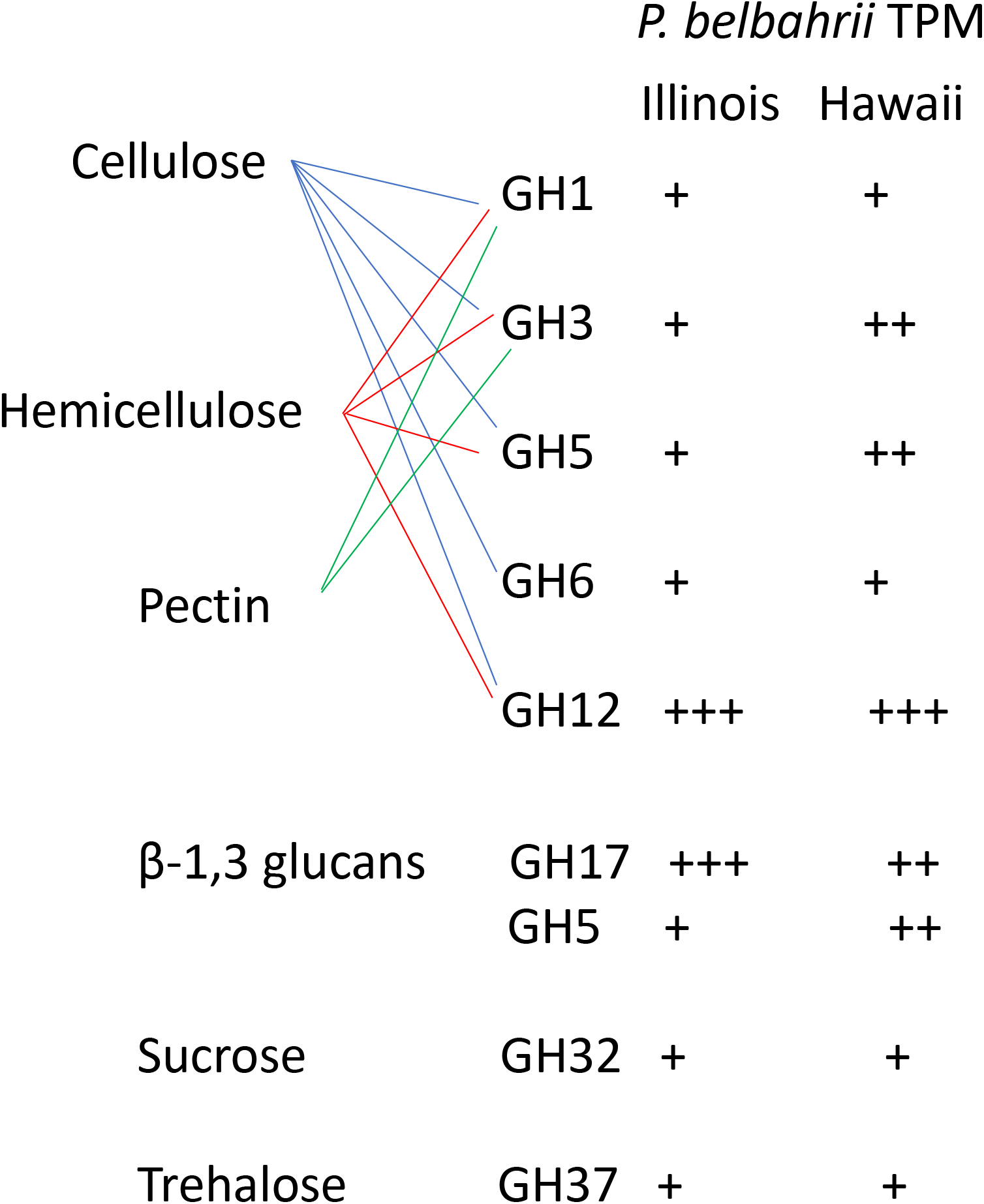
Possible substrates utilized by *P*. *belbarhrii* GH (glycoside hydrolase) enzymes. Lines between the substrate and GH class indicates likely hydrolysis of substrate; ‘+’ depicts the relative level of mRNA encoding a certain class of GH in the Illinois or Hawaii transcriptomes.

PEBEL_30158 and PEBEL_30235 encode GH12 enzymes, which have been characterized as virulence factors in a few *Phytophthora* species (Ma *et al.* 2015). BLAST analysis determined that PEBEL_30158 and PEBEL_30235 proteins were distantly related to the *Phytophthora* GH12 proteins (28-35% identity). Most GH12 proteins contain two conserved glutamate residues that contribute to catalysis; site directed mutagenesis of either of these residues in recombinant *Phytophthora sojae* GH12 enzyme abolished xyloglucan hydrolysis (Ma *et al.* 2015). Surprisingly, the PEBEL_30158 and PEBEL_30235 enzymes do not contain the conserved glutamate residues present in GH12 enzymes in many fungi and oomycetes (Supplementary File 10), and may indicate different amino acids critical for catalysis.

PEBEL_04311 transcripts, encoding a GH17 enzyme, were highly expressed in the Illinois transcriptomes and moderately expressed in the Hawaii transcriptomes. GH17 enzymes hydrolyze β-1,3-glucans (Pitson *et al.* 1997; Minic 2008; de Sousa Lima *et al.* 2012). GH17 transcripts were expressed by *Phytophthora parasitica* 48-60 hours after inoculation onto lupin roots (Blackman *et al.* 2015). It has been hypothesized that GH17 enzymes degrade the β-1,3-glucans synthesized by plants to make cell wall papillae at infection sites (Blackman *et al.* 2015). Papillae synthesis is part of the plant defense response and these specialized structures might impede the progress of pathogen growth (Underwood 2012). PEBEL_04261 transcripts, which also contain a GH17 motif, were well expressed in the Hawaii transcriptomes but not in the Illinois transcriptomes. The PEBEL_04261 transcript also encodes a region in the C-terminus with high homology to a herpesvirus protein (E-value: 6.91e-09 using CD-Search on the NCBI Conserved Domain Database); this region of 395 amino acids contains 31% proline residues and 28% serine residues. The herpesvirus protein is named UL36, an essential inner tegument protein that is 3164 amino acid residues and functions as a reversible, multivalent cross-linker protein between the capsid and the tegument during virus assembly (Newcomb and Brown 2010; Cardone *et al.* 2012; Schipke *et al.* 2012). This proline-serine rich region of PEBEL_04261 might serve as a protein cross-linker for the GH17 activity in the N-terminus of the protein, or might have an altogether different role. In any case, GH17 enzyme activity likely impedes the ability of basil to build physical defense structures against *P. belbahrii*.

A number of *P. belbahrii* genes encoding effectors, as well as a gene encoding an NLP, were expressed in the basil leaves after inoculation; these virulence genes were expressed at levels similar to the housekeeping genes listed in Table 4. None of the expressed effector genes in this dataset have sequence homology with avirulence genes from *Phytophthora*. There are 14 effector genes in *P. belbahrii* that have the exact RxLR motif (Thines *et al.* 2020), four of which were expressed in the Illinois and Hawaii transcriptomes (PEBEL_01944, PEBEL_03285, PEBEL_06440, PEBEL_07289). These effectors should be further characterized in a heterologous system to identify their phytotoxicity (or suppression of cell death) and localization properties (Liu *et al.* 2018).

There are ten genes in *P. belbahrii* that encode an elicitin like protein (Thines *et al.* 2020); four genes encode nearly identical proteins while two genes encode the same protein. However, only seven of the elicitin like proteins encoded by these genes have the conserved six cysteine residues that are found in most elicitin and elicitin-like (ELL) proteins (Jiang *et al.* 2006). PEBEL_03222 encoded protein is similar to elicitin-like protein SOL11B of *P. sojae*; a low level of *sol11b* transcripts were detected in soybean tissue infected by *P*. *sojae*, but other elicitin transcripts such as *soj1a1* and *soj5* were more prevalent in these infected tissues (Jiang *et al.* 2006). PEBEL_03222 is also similar (E value of 8e ^-45^ with BLASTP at NCBI) to an elicitin-like gene in *Plasmopara viticola* that is well expressed in sporangia, germinated zoospores and grapevine leaves 48 h after inoculation (Mestre *et al.* 2012). It is hypothesized that elicitins and ELLs sequester sterols for growth (Ponchet *et al.* 1999). There are three other *P. belbahrii* genes that encode nearly the same transcript as PEBEL_03222 (PEBEL_06439, PEBEL_06444, and PEBEL_06445); we did not explore which *P. belbahrii* gene is synthesizing this ELL transcript.

A disulfide isomerase gene expressed by *P. belbahrii* (PEBEL_06679) might be contributing to infection. The EffectorP 3.0 software program (Sperschneider *et al.* 2016; Sperschneider *et al.* 2018) predicted that PEBEL_06679 is a cytoplasmic effector. *Phytophthora parasitica* overexpressing a disulfide isomerase-GFP fusion protein made more haustoria-like structures after inoculation onto *N*. *benthamiana* and displayed enhanced virulence over the wild type pathogen (Meng *et al.* 2015). The *P*. *parasitica* disulfide isomerase possibly contributes to protein folding in the endoplasmic reticulum and elevated expression appears to facilitate virulence; more study is needed to determine the role of the *P*. *belbahrii* disulfide isomerase during infection of basil.

A single cyclophilin gene was well expressed in the Hawaii and Illinois *P. belbahrii* transcriptomes. There are 16 and 28 cyclophilin genes in *Peronospora effusa* and *Peronospora tabacina*, respectively (Zhang *et al.* 2020). There are likely a similar number of cyclophilin genes in *P*. *belbahrii*, but only one gene, PEBEL_06536, encodes a secreted protein. PEBEL_06536 is part of the oomcCYP00-i orthogroup; signal peptides or transmembrane domains were often located in the proteins that were part of oomcCYP00-i (Zhang *et al.* 2020). A cyclophilin has peptidyl-prolyl *cis-trans* isomerase activity which catalyzes the trans-cis isomerization of peptide bonds with proline residues (Zhang *et al.* 2020). While it is clear that most oomycetes have several cyclophilin genes, no function during pathogenesis has been described.

There are other proteins secreted by *P*. *belbahrii* that likely contribute significantly to successful pathogenesis in basil, but this study has focused on genes encoding cell wall digestive enzymes and a handful of virulence proteins. Based on this study, *P*. *belbahrii* invests significant resources into synthesizing many GH12 and GH17 transcripts. Pathogens mutated in a single glycoside hydrolase enzyme exhibited reduced virulence on their host (Rajeshwari *et al.* 2005; Brito *et al.* 2006; Ma *et al.* 2015). This indicates that novel strategies to inhibit GH12 and GH17 activities in *P*. *belbahrii* in the initial stages of infection could result in reduced downy mildew disease in basil. Alternatively, a better understanding of the mechanisms of the *P*. *belbahrii* virulence proteins could lead to control strategies. This fundamental knowledge of the pathogenesis employed by *P*. *belbahrii* is important to investigate since it is highly possible that current resistance genes employed by basil could be overcome by the pathogen over time.

## Acknowledgments

We thank Liis Kolberg for assistance with the use of g:Profiler, Mark Doehring for excellent technical assistance and Shyam Kandel for comments on the draft manuscript. The mention of firm names or trade products does not imply that they are endorsed or recommended by the USDA over other firms or similar products not mentioned. USDA is an equal opportunity provider and employer.

## Funding

Funding for this work was provided to EJ using USDA ARS in-house project 5010-22410-017-00-D, to HSK using USDA ARS in-house project 5010-42000-053-00D and NIFA HATCH project (accession number 1020611) to MT.

## Author contributions

EJ conceived the idea. IG, HK, and OT contributed expertise in bioinformatic analysis. MT contributed expertise in RNA extraction and cDNA synthesis. All authors contributed to the manuscript.

## Conflict of interest

All authors certify that there is no conflict of interest with any financial organization regarding the material discussed in the manuscript.

## Supplementary files

File 1. Mapping of transcriptome reads to genomes

File 2. Representative example of basil leaves infected by *P*. *belbahrii* at 6 dai

File 3. Results of differential gene expression in basil

File 4. Additional PCA of transcriptomes

File 5. Results of enrichment analysis for basil differentially expressed genes

File 6. Results of mapping the *DMR6-like* genes in the basil genome

File 7. All sequences used in Figure 3

File 8. All *P*. *belbahrii* gene sequences used for expression analysis

File 9. Expression results of *P*. *belbahrii* genes

File 10. Multisequence alignment of several GH12 enzymes

File 11. Collection of *P*. *belbahrii* ST and SWEET proteins

## Notes

### Competing Interest Statement

The authors have declared no competing interest.

