## Supplementary material for "Dual transcriptional analysis of *Peronospora belbahrii* and *Ocimum basilicum* in susceptible interactions": SUPP FILE2

### Slide 1
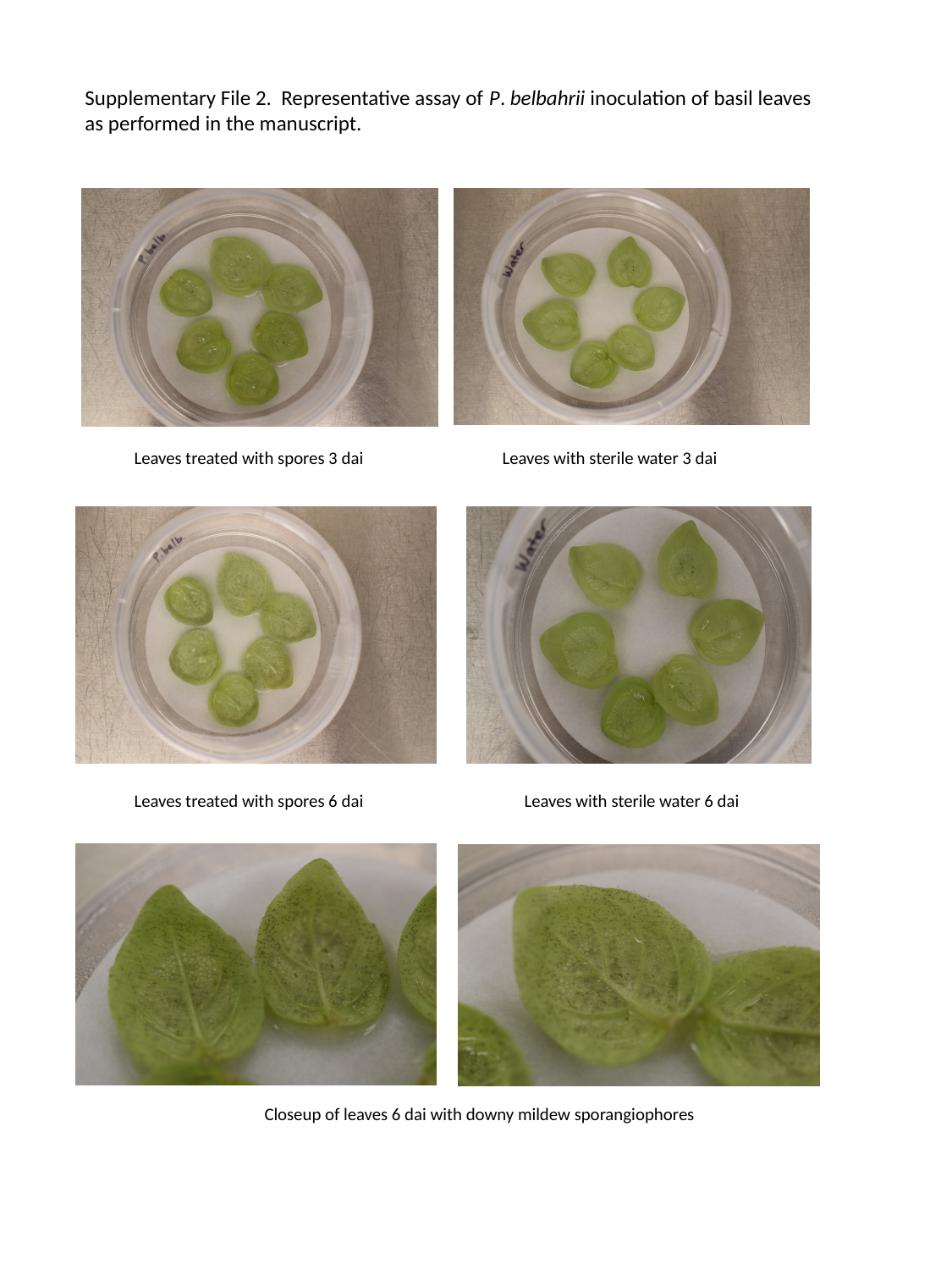

Supplementary File 2. Representative assay of P. belbahrii inoculation of basil leaves
as performed in the manuscript.
Leaves treated with spores 3 dai
Leaves with sterile water 3 dai
Leaves with sterile water 6 dai
Leaves treated with spores 6 dai
Closeup of leaves 6 dai with downy mildew sporangiophores
