## Supplementary figures and images for "Dual transcriptional analysis of *Peronospora belbahrii* and *Ocimum basilicum* in susceptible interactions"

### SUPP FILE4

#### Slide 1
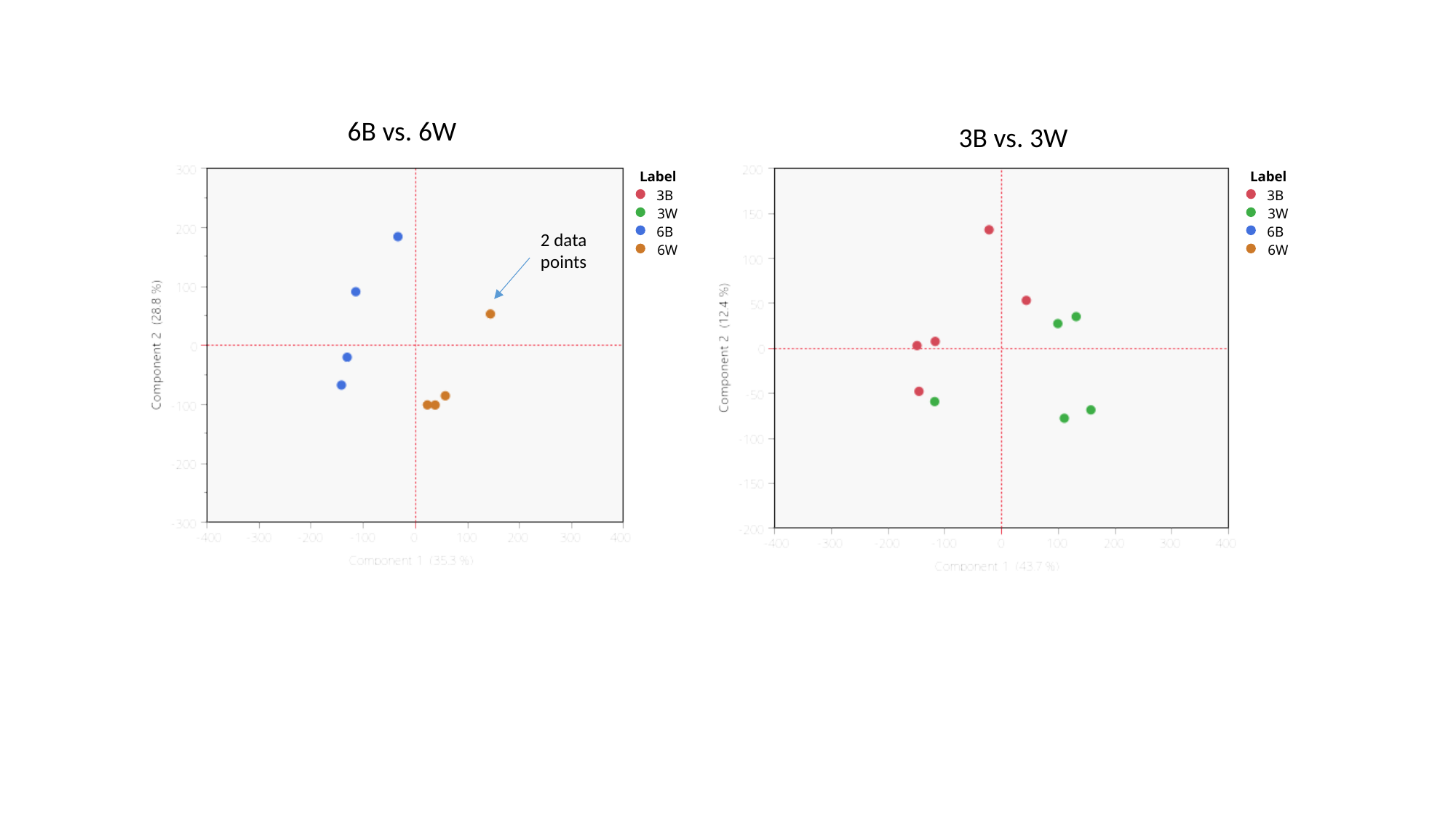

6B vs. 6W
3B vs. 3W
Label
3B
3W
6B
6W
Label
3B
3W
6B
6W
2 data points

#### Slide 2
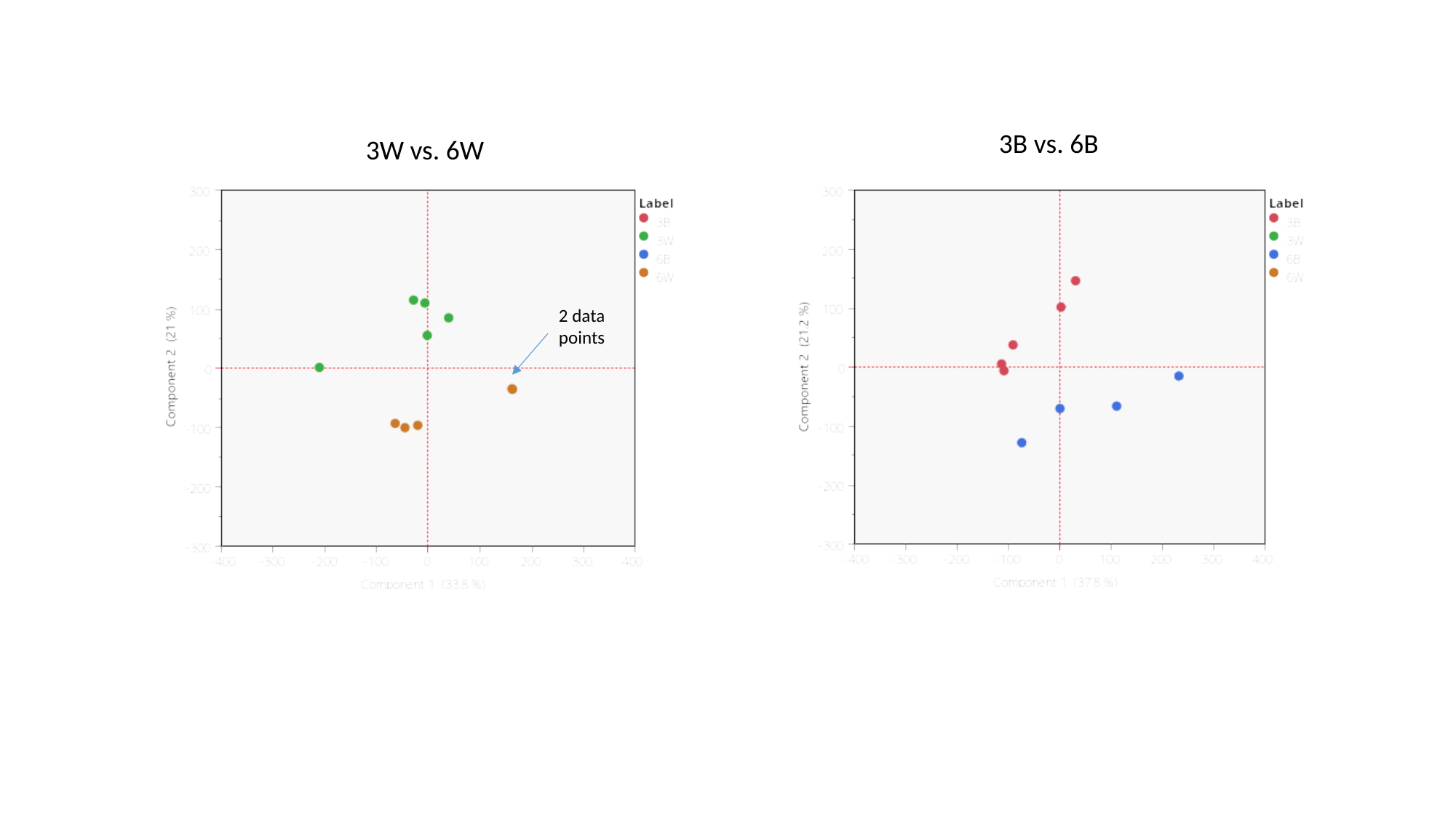

3B vs. 6B
3W vs. 6W
2 data points
