## Supplementary material for "Dual transcriptional analysis of *Peronospora belbahrii* and *Ocimum basilicum* in susceptible interactions": SUPP FILE10

```
Supplemental file 10. CLUSTAL Omega (1.2.4) multiple sequence alignment of several oomycete GH12 enzymes.  The coding for the organisms is at the end of the alignment.


PEBEL_30235           -MKLLLATAAIAACLATPLTSSLTTTISC-NVPS--ISENVLIGPNHKDYEEFCNQH---	53
PEBEL_30158           MMKLFIAAAALAVSFTTALNTRLA------AEP----SSSDLYAKRYENNKNYCGEV---	47
HpaG808599            -MKILVPAIPLALVA----------------------------TSSLTAAQRFCGQY---	28
HpaG814377            -MKILVPAIPLALVV----------------------------TFSLTAAQRLCGQY---	28
P_capsici_7335        -MKFLLPATLALA------------------------------AVASTNAADFCTFNNDD	29
HpaG805382            -MKVASSTAMMTA------------------------------AFSVAYAADTCDQW---	26
P_sojae_138787        -MKVLFATALTAA------------------------------AVAVASAAEFCDQW---	26
P_capsici_99469       -MKVFFAAALTAA------------------------------AVSVASAADFCDQW---	26
PPTG_16566            -MKVFFAAALTAA------------------------------AMSAAYAEEFCDQW---	26
PITG_06962            -MKVFFAAALTAA------------------------------AMSATYAEEFCDQW---	26
PPTG_19378            -MKSFLIAIV-IA------------------------------VLLPVSAADFCAQW---	25
P_sojae_144813        -MKSFLQLVVVVA------------------------------ALLSVSTADFCSQW---	26
P_capsici_111741      -MKGFFTGVIAAA------------------------------AFAVASAGEYCGQW---	26
P_sojae_109681        -MKGFFAGVVAAA------------------------------TLAVASAGDYCGQW---	26
PPTG_19377            -MKGLLAGTIAAA------------------------------TFAVASAGEYCGQW---	26
PPTG_16567            -MIMLYAATVTAL------------------------------TLAAVYADDFCDQW---	26
P_sojae_119627        -MKVAFATAMAAA------------------------------ALAAAYADDFCDQW---	26
P_capsici_98553       -MKVLFTVALTAT------------------------------TLATAHAADFCDQW---	26
P_sojae_140303        ---MKFSLTAVAA-----------------------------FAASAVSADKMCGDW---	25
PPTG_16278            -MKFLFAVVTLSA-----------------------------FAASTYAEKAMCGDW---	27
PPTG_19219            -MKLPIAILA---VG---------------------------LSTTAITGTEFCGEW---	26
PITG_08944            -MKL--LIPATLSLV---------------------------ALASTVDGQKFCGRD---	27
PPTG_16271            -MKF--LIPATLALA---------------------------AVASSADAQKFCGRD---	27
P_capsici_15451       -MKF--LIPATLAVA---------------------------ALVSSTEAQQYCGRD---	27
P_sojae_140289        -MKF--LLPATVAVA---------------------------ALASSASAQQFCGRD---	27
P_sojae_109280        -MKF--LLPATVTIA---------------------------ALASSASAQQFCGRD---	27
PPTG_16267            -MKF--LIPATLAVA---------------------------AVASSANAQEYCGRN---	27
P_capsici_106898      -MKF--LLPVTIAVA---------------------------AVA-STNAQEYCGRN---	26
P_sojae_140300        -MKLSVAIAFAAAAV---------------------------AA-PPAVAKEFCDLW---	28
PITG_16991            -MKFTVVIATAATAV---------------------------AALPVFAEEEFCGQW---	29
PPTG_16273            ---MKISIAITVAAV---------------------------AASSVFAEEEFCGQW---	27
P_capsici_107440      -----MKLSVAIAAV---------------------------AASAVAAEQEFCGQW---	25
PITG_13560            -MKLPLVIAATAATV---------------------------AAFPAFAKDEFCGQY---	29
P_sojae_109281        -MKLSAAVLA---LA---------------------------AAASATDAKEFCGRN---	26
PPTG_16272            -MKLSFAALA---LA---------------------------AAASSSNAQEFCGRD---	26
P_capsici_107334      -MKLSAAALA---LA---------------------------ATASSATAQEFCGRN---	26
PPTG_19220            -MKFFAIAAAIAVAVGSTSAKDITQQTTQSSDPSTQQTTQNEGSSTTGAVEDFCGQW---	56
P_capsici_15453       -MKLFAIIAALAVASTSASTTQHQQTTQDS------SSAEGSASFATEATGDFCGQW---	50
P_capsici_107122      -MKLFTIIAALAVASTSASTTQHQQTTQDS------SSAEGSASFATEATGDFCGQW---	50
P_capsici_15454       -MKLFTIIAALAVASTSASTTQHQQTTQDS------SSAEGSASFATEATGDFCGQW---	50
P_sojae_140302        -MKSSAVLAASALAI---------------------------AASPAFAQEEFCGQW---	29
PITG_16992            -MKLAVALATVAF-------------------------------VAASVKSELCGEQ---	25
P_sojae_140301        -MQISTVLTASALAL---------------------------AASPSFAEKEFCGEK---	29
P_capsici_107461      -MKFSTIFAAAAV------------------------------TASSVSAETFCGEQ---	26
PITG_14060            -MKLSAALTTIAI-------------------------------ATASAETEFCGEK---	25
                                                                           *

PEBEL_30235           -DMAERGNYFFVTNTHVEKYWTMFDSHCTAVT--DYSNEKIGLEIFF-DIKD-GNSMVRS	108
PEBEL_30158           -ETIPRESFIVFQNRNGADKR---VGWCIAVT--SSSKNDVSWYAQLNADDSLKPSPVNL	101
HpaG808599            -DLKVVPPYTVYNNLWGQADDS-NGTQCTEVT--RISDESIAWTTDF-KWAG-SRYQVKS	82
HpaG814377            -DEIVLPPYTVYNNLWGQGDDP-NGIQCTEAT--RILDKAIAWTTDF-NWAG-NPNQAKS	82
P_capsici_7335        SSKAVAGAYTVYNNVGESNV---NAQQCTGVD--SSEGTSIAWHTSY-NTTG-DDALSKS	82
HpaG805382            -GQTKSGNYIIYNNLWGSSAATGGGKQCTGLD--SGSGDTVAWHTVW-SWMG-GDTSVKS	81
P_sojae_138787        -GQAKSGNYIIYNNLWGSSAANPGGKQCTALD--SGSGDSVAWHTTW-SWQG-GDKSVKS	81
P_capsici_99469       -GQVKSGNYIIYNNLWGSSAATGGGKQCTGLA--SGSGDTVSWHTKW-SWQG-GDKSVKS	81
PPTG_16566            -GKTTSGNYIIYNNLWGSSAATGGGKQCTSLN--SGSGDTVSWQTTW-SWQG-GDKEVKS	81
PITG_06962            -GKTTSGNYIIYNNLWGSSAATGGSKQCTSLN--SGSGDTVAWQTTW-SWQG-GDKEVKS	81
PPTG_19378            -RLSKAGKYIIYNNLWNQNTATS-GSQCTGVD--KVSGSTVAWHTSY-SWAG-APTQVKS	79
P_sojae_144813        -RLSKAGKYVIYNNLWNKNAAAS-GSQCTGVD--KISGSTIAWHTSY-TWTGGAATEVKS	81
P_capsici_111741      -DWAKSSQYTVYNNLWNKKAAAS-GSQCTGVD--KISGSTIAWHTSY-TWTGGAATEVKS	81
P_sojae_109681        -DWAKSTNYIVYNNLWNKNAAAS-GSQCTGVD--KISGSTIAWHTSY-TWTGGAATEVKS	81
PPTG_19377            -DWAKSTQYTVYNNLWNKNAAAS-GSQCTGVD--KISGSTIGWHTSY-TWTGGAATEVKS	81
PPTG_16567            -GTATTDNYILYNNLWGESYATS-GSQCTGLD--SSSGSTISWHTNW-TWAG-ASSNVKS	80
P_sojae_119627        -GTTTTDNYIIYNNLWGESYATS-GSQCTGLD--SSSGSTVAWHTNW-TWTG-ASSNVKS	80
P_capsici_98553       -GTATTDNYIIYNNLWGESYATS-GSQCTGLD--SNSGSTVAWHTNW-TWVG-ASSSVKS	80
P_sojae_140303        -DTAVVGDFTLYNNLWGKENDPGTGHQCTTLLNSSADNSTLGWSTEY-KWGG-ITWNVKG	82
PPTG_16278            -DTAVVGDYTLYNNLWGKDNDPGTGHQCTTLLSTSADNSTIGWTTNY-KWGG-ITWNVKG	84
PPTG_19219            -NSTETSDYILYNNLWGAFDDP-NGSQCTGLD--SVDGSTIAWHMRY-NWAG-TPYQVKS	80
PITG_08944            -DRKVVGDYTVYNNLWGEDNDK-SGAQCTEVT--GSTSSSVSWKTSF-NWAG-DKWQVKS	81
PPTG_16271            -DLKVVGDYTVYNNLWGEDNDK-TGGQCTEVT--GSTSSSVSWKTSF-NWAG-DNWQVKS	81
P_capsici_15451       -DFKVVGDYTVYNNLWGEDNDK-TGGQCTEVT--GSSGSSVSWKTKF-NWAG-DNWQ---	78
P_sojae_140289        -DNKVVGDYTVYNNLWAEENDP-KGGQCT-----------------F-NWTG-DSWQVKS	66
P_sojae_109280        -DNKVVGDYTVYNNLWAEENDP-KGGQCTEVT--GSTSSSVSWQTSF-NWAG-DNWQVKS	81
PPTG_16267            -DLKVVGDYTVYNNLWGQDNDK-TGKQCTEVT--GSTSTSVSWQTSF-NWAG-DSWQVKS	81
P_capsici_106898      -DLKVVGDYTVYNNLWGQDNDK-TGKQCTEVT--GSTSTAVSWQTSF-NWAG-DSWQVKS	80
P_sojae_140300        -GSAQTDDYTVYNNLWGAHDDP-KGGQCTELD--SVDGSNIAWHTSF-NWNG-SSWQVKS	82
PITG_16991            -SSTQTADYIIFNNLWGQDNDK-AGGQCTGLD--SVDGSTIAWHTSF-NWAG-DNWQVKS	83
PPTG_16273            -NSTQTDDYTIYNNLWGQDNDK-AGGQCTGLD--SVDGSTIAWHTSF-NWAG-DNWQVKS	81
P_capsici_107440      -NSTQTDDYTIYNNLWGMDNDK-AGGQCTGLD--SVDGSTIAWHTSF-NWAG-DNWQVKS	79
PITG_13560            -DSIQAGDYIVHNNLWGQDNDK-AGGQCTGLD--SIDGSTIAWHMSFPSWVG-DNWQVKS	84
P_sojae_109281        -DMQFLGDYTVYNNLWGEDNDP-EGGQCTEVD--GSSDSEVSWHTHF-NWAG-DNWQVKS	80
PPTG_16272            -DFKVVGDYTVYNNLWGQDNDK-AGGQCTTVD--GSSGSEIAWHTSF-NWAG-DNWQVKS	80
P_capsici_107334      -DLQVVSDYTVYNNLWGQDNDK-TGGQCTTVD--SSSGSEIAWHTSF-NWAG-DNWQVKS	80
PPTG_19220            -NLTETEHYTLYNNLWGAHNDT-KGTQCTGLD--SVEGSTIAWHTSY-NWKG-SKLQVKS	110
P_capsici_15453       -NTTEAADYTLYNNLWGQSNDT-AGTQCTGLD--SVDGAHISWHTSY-NWKG-SKFQVKS	104
P_capsici_107122      -NTTEAADYTLYNNLWGQSNDT-AGTQCTGLD--SVDGAHISWHTSY-NWKG-SKFQVKS	104
P_capsici_15454       -NTTEAADYTLYNNLWGQSNDT-AGTQCTGLD--SVDGAHISWHTSY-NWKG-SKFQVKS	104
P_sojae_140302        -NLTKTDDYILYNNLWGAFDDP-TGTQCTGLD--SVKGSTVAWHTSF-GWSG-SKTQVKS	83
PITG_16992            -NKTETDNYILYNNLWGAFDDP-RGHQCRGLD--SVNDSTIAWHTSF-TWNG-TAWQVKS	79
P_sojae_140301        -NLTQTDDYILYNNLWGAWDDP-KGWQCTGLD--SVDGSTISWHTRF-SWNG-TAWQVKS	83
P_capsici_107461      -NKTETADYILYNNLWGAYDDP-KGHQCTSLD--SVDGSSISWSTNF-SWNG-TAWQVKS	80
PITG_14060            -NKTETADYVLYYNLWGAFDDP-NGHQCTGLD--SVDGSTISWHTSF-SWNG-TAWQVKS	79
                              : .  *             *                       .

PEBEL_30235           YSHVDITFKPVLPLYISSIPFSMEYRIKHDDSASLNVYARCLFYANVH-----TTYSLTV	163
PEBEL_30158           TANVALKYNTIVLYVVTSIKTMMKYEYIAG-PGVIANVAYNLEMNTEFQTNTGVQYFIRV	160
HpaG808599            FANAALTFTPVKMSQVASMPTVIEYKYESVDDTLITNVAYDMLLSPHP-EGE-FTYELMV	140
HpaG814377            FAIVALSFTPVQMSQVTSMPTEIEYEYESVDKTLVANVAYDMFLSSSS-DGETYTYEVKV	141
P_capsici_7335        FVVASLQMNQTKLAELQSIPTEVTYTYTSQ-GEMISNAVYDMYTTS---YGH-PQYKVMV	137
HpaG805382            YANAALTFDAVPLTEVDSIPSSIEFKVKYS-GKAVANVAYDLFTSSTA-EGE-KKFEIMI	138
P_sojae_138787        FANAALEFDPVPLSEVKSIPSTMAYTVKSK-GKAVTDVAYDLFTSSTA-KGE-KEFEIMI	138
P_capsici_99469       FANAALEFDPVPLTEVKSIPSTMSYTVKYS-GNVVADVAYDLFTSSTA-KGE-KEFEIMI	138
PPTG_16566            YANAALEFEPVPLTEVKSIPSTMSYKVKYS-GKVVADVAYDLFTSSTA-KGE-KEFEIMI	138
PITG_06962            YANAALEFEPVPLTEVKSIPSTMSYKVKYS-GKVVADVAYDLFTSSTA-KGE-KEFEIMI	138
PPTG_19378            YSNAALVFTKKQIKNIKTIPTTMKYSYSYS-GTLIADVAYDLFTSSTA-SGS-NEYEIMI	136
P_sojae_144813        YSNAALVFSKKQIKNIKSIPTKMKYSYSHSSGTFVADVSYDLFTSSTA-SGS-NEYEIMI	139
P_capsici_111741      YSNAALIFTPKQVKSIKSIPTTMKYSYSYSAGKFVADVAYDLFTSSTA-TGS-NEYEIMI	139
P_sojae_109681        YSNAALVFSKKQIKNIKSIPTKMKYSYSHSSGTFVADVSYDLFTSSTA-SGS-NEYEIMI	139
PPTG_19377            YSNAALIFSPKQIKNIKTIPTKMKYSYSHSSGTFVADVSYDLFTSSTA-TGK-NEYEIMI	139
PPTG_16567            YANAALQFDAVQLSSISSIPTTMDYSLDYS-DTIVADVSYDLFTSSTS-NGD-NEFEVMI	137
P_sojae_119627        FANAALQFDAVQLSSVSSIPTTMEYSLEYS-GNIAADVSYDLFTASTS-TGD-NEFEIMI	137
P_capsici_98553       YANAALQFDAVQLSSVSSIPSTIEYSLDYS-GTIVADVSYDLFTSSTS-DGD-NEFEIMI	137
P_sojae_140303        YPNVGLHFKPIELANVESIPTSINYTYEYS-PTTTANVAYDLFTSSTP-DGD-FEYEIMV	139
PPTG_16278            YPNVGLHFKPLELANVESIPTTINYTYEHE-PTTTANVAYDLFTSSTP-DGD-FEFEIMV	141
PPTG_19219            FPNAALKFDPVQLLNVKSIPATIEYEFEYS-ENTIANVAFDLFTRSTI-DGA-VEYEVMV	137
PITG_08944            FANAALKFQPKQISAIKSIPTSIEYTYTYG-DHIITNVAYDLFTSSSP-TGE-TEYELMV	138
PPTG_16271            YANAALKFQPKQVSAISSIPTSIDYTYTFD-GHIIANVAYDLFTSSSA-NGE-IEYELMV	138
P_capsici_15451       -----------QVSAISSIPTSIDYTYTFD-GHIIANVAYDLFTSSSA-NGE-IEYELMV	124
P_sojae_140289        FANAALN---------------TAYNYTYD-GKIIANVAYDMFTSSTE-SGD-IEYELMV	108
P_sojae_109280        FANAALKFTPKQVSTVTSIPTSIAYKYTYD-GNIIANVAYDLFTSSTP-TGD-VEYELMV	138
PPTG_16267            FANAALKFQPKQVSAITSMPTTMKYEYTYD-GNIIANVAYDLFTSSSE-SGE-IEYELMV	138
P_capsici_106898      FANAALKFEPKQVSAITSMPTTMQYEYTYD-GNIIANVAYDLFTSSSA-NGE-IEYELMV	137
P_sojae_140300        FANAALKFEPVQLANISSIPSTIEYDYKYD-GKIITNVAYDLFTSATA-NGT-VEYELMV	139
PITG_16991            FANAALKFAPVQISSVASIPSTIEYDYKYD-GNIITNVAYDLFTSATA-NGT-VEYELMV	140
PPTG_16273            FANAALKFNPVQIANVASIPSTIEYDYKYD-GNIITNVAYDLFTSATA-NGT-VEYELMV	138
P_capsici_107440      FANAALKFDPVQLANVSSIPSTIEYDYKYD-GDIITNVAYDLFTSATA-KGN-VEYELMV	136
PITG_13560            YANAALKFNLVQLSSVKSIPTTMEYTYKYD-GNIITNVAYDLFTSPSI-GGE-TAYELMV	141
P_sojae_109281        YANAALKFDPVQVANVKSIPTTMSYSYTYD-GNIITNVAYDLFTSPTI-GGE-TAYELMV	137
PPTG_16272            YANAALKFDPVQISNVTSIPTTMEYTYKYD-GNIITNVAYDLFTSPSI-GGE-TAYELMV	137
P_capsici_107334      YANAALKFDPVQISNVTSIPTTMEYTYKYD-GNIITNVAYDLFTSPTI-GGE-TAYELMV	137
PPTG_19220            FANVALKFDKVPLSTVKSIPSTIKYDFKSK-GKVIANVAYDLFTTSTP-DGD-AEYEIMV	167
P_capsici_15453       YANAALKFDKVPLSKVKSIPSTIQYDFKSR-GKVVANIAYDLFTTSTP-DGD-AEYEIMV	161
P_capsici_107122      YANAALKFDKVPLSKVKSIPSTIQYDFKSK-GKVVANIAYDLFTTSTP-DGD-AEYEIMV	161
P_capsici_15454       YANAALKFDKVPLSKVKSIPSTIQYDFKSK-GKVVANIAYDLFTTSTP-DGD-AEYEIMV	161
P_sojae_140302        FANVALQFDHLPISEVTSIPSTIHFKYDYE-ESLIANVAFDLFTSSTP-DGD-AEYEIMV	140
PITG_16992            FANAALKFDHVPIANVTSIPSVIEFNFDYE-GKLVANVAFDMFTASTL-GGT-AEYEVMV	136
P_sojae_140301        FANAALQFDHVPIADVTSIPSTIEYDFQYE-GKTVANVAYDLFTTSKI-GGD-AEYEVMV	140
P_capsici_107461      FANAALKFDHVPIANVTSIPSTIEFDFAYE-GKLVANVAFDMFTASKL-GGD-AEYEVMV	137
PITG_14060            FANAALKLDHVPIANK----------------KLVANVAFDMFTTSSL-GGN-AEYEVMV	121
                                                               :             : : :

PEBEL_30235           LFELVGGMKNLINLK----KLNDDVKIGEDSYELYDGIFGNFEGYIFVANKGNRRLNIDF	219
PEBEL_30158           WINALGGNIPYGDTV----PGK-GLIVSNVAYDLYIGNHNGIRYITYVATQPVTHFDDDL	215
HpaG808599            WLATFGGAAPLARSYDPMVPVKANVTVAGVNFNLYEGMNGNVTVLTYLATGSINRFSGNL	200
HpaG814377            WLTTFGIIAPFV--EDPMNPILANVTVAGVNFKLYRGMDSNVTVFTYVATANINRFNGDF	199
P_capsici_7335        WLASYGGAKLKSTTG---LPIK-TATVGGVEFDLYQGYYQNMGVYTFVAKTPLTTFKGDL	193
HpaG805382            WMAAIGGAGPISSTG---KPID-TAKIAGVEWSVFSGPNGQMVVYSFVASKPSNNFEGDL	194
P_sojae_138787        WLAALGGAGAISSTG---KPIA-STTIAGTEWSVYKGPNGSMMVYSFVASKQVENFEGDL	194
P_capsici_99469       WLAAIGGAGPISSTG---KAID-TTTIAGTEWSVYKGPNGQMMVYSFVASKQVENFEGDL	194
PPTG_16566            WLAAIGGAGPISSTG---KAID-TTTIAGNEWSVYSGPNGQMMVYSFVASKQVENFEGDL	194
PITG_06962            WLAAIGGAGPISSTG---KALA-TTTIAGTEWSVYSGPNGQMMVYSFVAFKQVENFEGDL	194
PPTG_19378            WLAAYGGAGPISSTG---KAIA-TVTINSNSFKLYKGPNGSTTVYSFVATKTITNFSADL	192
P_sojae_144813        WLAAYGGAGPISSTG---KAIA-TVTIGSNSFK---------------------------	168
P_capsici_111741      WLAAYGGAGPISSTG---KAIA-TVKIGSNSFKLYKGPNGSTTVFSFVATKTITSFSADL	195
P_sojae_109681        WLAAYGGAGPISSTG---KAIA-TVTIGSNSFKLYKGPNGSTTVFSFVATKTITNFSADL	195
PPTG_19377            WLAAYGGAGPISSTG---KAIA-TVTIGSNSFKLYKGPNGSTTVFSFVATKTITNFTADL	195
PPTG_16567            WLAALGGAGPISSTG---SAVA-TTTIADTEFSLYSGANGDTTVYSFVASDTVKSFSGDL	193
P_sojae_119627        WLAALGGAGPISSTG---SAVA-TTTIADTSFSLYTGANGDTTVYSFVASDTVKSFSGDL	193
P_capsici_98553       WLAAIGGAGPISSTG---SAVA-TTTIANTEWSLYSGANGDTTVYSFVASDTVKSFSGDL	193
P_sojae_140303        WLAAIGTAWPLASEG---KNIK-NITVSGVDFMLNHGVNGNMTVFSYVATQLTENFSGDL	195
PPTG_16278            WLAAIGTAWPLSSEG---KNIK-NITVSGVEFMLNHGVNGNMTVFSYVASKLTENFSGDL	197
PPTG_19219            WPAALGGALPLSTSG---KPIK-TTNIGDVDFTLYQGMNGNMTVLSYVPDKMITNFSTDL	193
PITG_08944            WLAALGEPWPLTDSG---KSIA-TVNIGGVVFELFQGMNKNVKVFSYVAKKTAYKFSADL	194
PPTG_16271            WLAALGGPWPLTDSG---KPIK-TVNIGGVDFDLFQGMNKKVKVFSYVAKKTAYKFSADL	194
P_capsici_15451       WLAALGGPWPLTDSG---KPIK-TVKIGGVEFDLFQGMNKKVKVFSYVAKKTAYKFSADL	180
P_sojae_140289        WLAALGDAWPLTSSG---QPIK-TFTAG-----------RKVKVFTYVAKQQATSFTADL	153
P_sojae_109280        WLSALGGAWPLTDSG---KPIK-TVTAGGVEFDLYQGFNKKVKVFSYVAKKSATSFTADL	194
PPTG_16267            WLAALGGAWPLTDSG---KPIK-SVTLGGVDFDLYQGMNKKVKVFSYVAKKTAKSFTADL	194
P_capsici_106898      WLAALGGAWPLTDSG---KPIK-SVTLGGVDFDLYQGMNKKVKVFSYVAKKTAKSFTADL	193
P_sojae_140300        WLAALGGAWPLTTSG---RPIK-AVNVGGTDFNLYQGKNGNTTVFSYVAVNSTTKFSADF	195
PITG_16991            WLAALGGAWPLTTTG---KPIK-EVKVGSVDFNLYQGKNGNTTVFSYVAVNTTKSFTADF	196
PPTG_16273            WLAALGGAWPLTTTG---KPIK-EVKVSNVDFNLYQGKNGNTTVFSYVAVNMTKSFSADF	194
P_capsici_107440      WLAALGGAWPLTTTG---KPIK-EVKISNVNFNLYQGKNGNTTVFSYVAVNTTKSFTADF	192
PITG_13560            WLAALGGAWPLTTTG---QPIK-SVKLGGVDFNLYQGWNNKTKVFTY-------------	184
P_sojae_109281        WLAALGGAWPLTTTG---QPIK-EVNLGGVDFNLYQGWNNKTKVFTYVAKQTTYTFEADL	193
PPTG_16272            WLAALGGAWPLTTTG---QPIK-SVTLGGVEFNLYQGWNNKTKVFTYVAKNMATSFSADL	193
P_capsici_107334      WLAALGGAWPLTSTG---QPIK-SVTLGGVDFNLYQGWNNKTKVFTYVAKNMATSFSADL	193
PPTG_19220            WLAAMGGAGPISATG---KPIKKGVKVGGVNFNLYHGKNGNMTVYSYVASKQTKSFSADF	224
P_capsici_15453       WLAALGGAGPISATG---KPIKKGVTVGGVKFDLWHGKNGNMTVYSYVAAKQTKSFKADF	218
P_capsici_107122      WLAALGGAGPISATG---KPIKKGVTVGGVKFDLWHGKNGNMTVYSYVAAKQTKSFKADF	218
P_capsici_15454       WLAALGGAGPISATG---KPIKKGVTVGGVKFDLWHGKNGNMTVYSYVAAKQTKSFKADF	218
P_sojae_140302        WLAAIGGAGPISSTG---SAVD-QVTVGGVDFSLYAGKNGNMTVYSFVASTMVNRYATDF	196
PITG_16992            WLQAIGGAGPLSNTE---KPIK-EVSIGGVNFSLIHGMNGNMTVFSYVAANTINSFSTDF	192
P_sojae_140301        WLEAIGGAGPLSNTG---SAIK-HVTVGGVDFSLYHGKNGNMTVYSYVAANAMNSFSADF	196
P_capsici_107461      WLQAIGGAGPLSNTG---KPIK-EVNVGGVDFSLYHGKNGNMTVFSYVAANTANNFSTDF	193
PITG_14060            WLQAIGGAGPLSNTG---RPVK-EVNVGGVDFSLYHGKNGNMTVFSYVAANTTNSFSTDF	177
                           *

PEBEL_30235           NGFFKALPKNSKINSNQYLYSLSVGSQVVQG-SGTVKFTGFSAAVNTSHTECGLRV----	274
PEBEL_30158           LVFINELKYNKFISDNDYLVKLDAGTRIIKGSSATFTVKKFQAAIKIESPAE--------	267
HpaG808599            QDFVEKLPNPKLL-DDQYLVKAETGTEPFQG-DAKLIVSKFSLEIIQKPSNAS-------	251
HpaG814377            KEFFTNLPDAKRL-NDQYLVDAMAGTEPSHG-KAKLIVSKYSLAIIS-------------	244
P_capsici_7335        KPFFSEIPADFT---EQALQAVYFGTEAFKGTNAKFSVSKLSVSLQ--------------	236
HpaG805382            MEFFDYLAKSQKFKMRQYLIKVECGMEPFIGKDVSLTVSKYSTVVKTGKSGSSTSSDSSA	254
P_sojae_138787        LEFFNYLVKEQGFKTSQFLIKVECGTEPFVGTDVTMTVSKYSAAVNT-SGGSSTPTQSGE	253
P_capsici_99469       MEFFNYLTKSQSFKTSQYLIKVECGTEPFVGTDVSMTVSKYSAAVNTGSGGSTTPTQSGD	254
PPTG_16566            MEFFNYLVKSQKFKTSQYLIKVECGTEPFVGNDVTMTVSKYSATVNTGSGGGSTPTQSGD	254
PITG_06962            MEFFNYLVKSHSFKTSQYLIKVECGTEPFIGKDVTMTVSKYSATVNTGSGGGSTPTQSGD	254
PPTG_19378            LDFFTYLVKTQAFPSSQYLTTLEAGTEPFTGSNAKMTVSSYSAAVN--------------	238
P_sojae_144813        ------------LPSTQYLTTLEAGTEPFTGSNAKMTVSSFSAAVN--------------	202
P_capsici_111741      QKFLSYLVKNQGLPSSQYLITVEAGTEPFVGTNAKMTVSSFSAAVN--------------	241
P_sojae_109681        QKFLSYLTKNQGLPSSQYLITLEAGTEPFVGTNAKMTVSSFSAAVN--------------	241
PPTG_19377            QKFLTYLVNSQGLPSSQYLITLEAGTEPFVGTNAKMTVSSYSAAVN--------------	241
PPTG_16567            MDFFTYLIDNESFSSSQYLNTVQAGTEPFTGTDVTLTVSSYSAVVNTGATSGTTSASSA-	252
P_sojae_119627        MDFFTYLIDNEGFSSSQYLNTVQAGTELFTGTDVTLTVSSYSAAVNMGASSGTTATTSTA	253
P_capsici_98553       MDFFTYLIENESFSTSQYLNTVQAGTEPFTGTDVTLTVSSYSAVVNTGASSGTVTTSSAT	253
P_sojae_140303        TEFIEALPQDVAPGAKQYLTKVQCGTESYHADRASMTVTSYTVAVNSVMQGDVGFSQN--	253
PPTG_16278            TEFIDNLPKDVAPDGKQYLTKVQCGTEAYHAENATMTVSAYTVAVNTLIQGDVGFSSSPS	257
PPTG_19219            KKFFDELPKSYAIARTQYLTHVQGGAEILVG-NGTLTVSKYQEAVHTTKHNSSKMTT---	249
PITG_08944            KKFFSELPKNNTLPQTQYLVKLEAGTEPFQGKNGEMIVKSYSSKVN--------------	240
PPTG_16271            KKFFNELPANNTLPQTQYLVKLEAGTEPFQGKNGEMIVKSYSSKVN--------------	240
P_capsici_15451       KKFFNELPANNTLPQEQFLVKLEAGTEPFQGKNGEMIVKSYSSKVN--------------	226
P_sojae_140289        KFFFDQHPTDNNLPTTQFLRKVEAGTEPFQGQNATMVVSSYSVQVK--------------	199
P_sojae_109280        KYFFDQLPADNNLPQTQYLQKVEAGTEPFQGKNAKMVVSSYSVQVK--------------	240
PPTG_16267            KQFFDELPADNTLPQTQYLQKVEAGTEPFQGKNAKFVVSTYSVQVK--------------	240
P_capsici_106898      KQFFDELPADNNLPQTQYLQKVEAGTEPFQGKNAKLVVSTYSVQVK--------------	239
P_sojae_140300        KQFFSELPADNTIAPTQFLTHVQAGTEPFQGQNATLTVSKYSVAVNSA------------	243
PITG_16991            KQFFDELPADNAIASTQYLTQVQAGTEPFQGKNATLTVSKYSAAVNS-------------	243
PPTG_16273            KQFFDELPADNTIAPTQYLTHVQAGTEPFQGKNATMTVCKYSAAVKSA------------	242
P_capsici_107440      KQFFDELPADNTIAPNQFLTHVQAGTEPFQGKNATLTVSKYSAAVHTA------------	240
PITG_13560            -QFFDELPADNTIETTQARTPL--------------------------------------	205
P_sojae_109281        RKFFDELPADNTIEATQYLTHVQAGTEPFQGKNATFTVHKYSAAVHTA------------	241
PPTG_16272            KQFFDELPADNTIETTQYLTHVQAGTEPFQGKNATMTVSKYSAAVHTV------------	241
P_capsici_107334      KQFFDELPADNTIEKTQYLTHVQAGTEPFQGKNATMTVTKYSAAVHAV------------	241
PPTG_19220            KQFFHKLPAKHTVSPKQYLTHVQAGTEPFVG-NGTMTVTEYAAAVHTV------------	271
P_capsici_15453       KQFFHNLPKKHAIAPTQILTHVQAGTEPFVG-NGTLTVSKYTAAVHTV------------	265
P_capsici_107122      KQFFHKLPKKHAIAPTQILTHVQAGTEPFVG-NGTLTVSKYTAAVHTV------------	265
P_capsici_15454       KQFFHNLPKKHAIAPTQILTHVQAGTEPFVG-NGTLTVSKYTAAVHTV------------	265
P_sojae_140302        KQFFDVLPRNLTIDPSQYLINVQAGTEPFVG-NGTLTVSKYSAAVNPAEYSQVQQQ----	251
PITG_16992            KQFFNELPAINTIAPDQYLINVQAGTEPFVG-NGTLTVTKYSTAVHTA------------	239
P_sojae_140301        KQFFDELPANNTIAPTQYLINVQAGTEPFVG-NGTLTVSKYSAAVHTAGAN---------	246
P_capsici_107461      KQFFDELPANNSIASEQYLVNVQAGTEPFVG-NGTLTVSKYTAAVHTA------------	240
PITG_14060            KQFFDELPSNNTIAPEQYLINVQAGTEPFVG-NGTLTVSRYSAAVNTAST----------	226
                                      :

PEBEL_30235           ------------------------------------------------------------	274
PEBEL_30158           ------------------------------------------------------------	267
HpaG808599            ------------------------------------------------------------	251
HpaG814377            ------------------------------------------------------------	244
P_capsici_7335        ------------------------------------------------------------	236
HpaG805382            KSDDDV-TQTDPTSSS--------------DNEVGGKAEAGDDLLS---ALGGTPSSDDK	296
P_sojae_138787        SSSSTEQTTTAPAASNTGSSAEQ-TTPAPAESNTGSSAEQT----------TPAPAGSDA	302
P_capsici_99469       SSSTGAQTTTAPSTGS---SNEQ-ST---TAPSTGSSSEQTPATSSTGSSSEQTPSTSST	307
PPTG_16566            SSSTGAQTTTAPSTGS---STEE-TP---STSTTGSSTEQNPSTPSTGSSTEETPATSST	307
PITG_06962            SSSTGAQTTTAPSTGS---STEE-TP---STSSTGSSIEQNPSTPSTGSSTEDTPSTGSS	307
PPTG_19378            ------------------------------------------------------------	238
P_sojae_144813        ------------------------------------------------------------	202
P_capsici_111741      ------------------------------------------------------------	241
P_sojae_109681        ------------------------------------------------------------	241
PPTG_19377            ------------------------------------------------------------	241
PPTG_16567            ------VTSTKELVASSSSTESETTV----------------------------PTATSS	278
P_sojae_119627        SSSTGTSVSTKELIASSSSTESDSQTAALT-------------------ATTATPSTDSS	294
P_capsici_98553       G-SSTTSTSTKELAASSSSTQTDTETTAPV-------------------A-----SSSRS	288
P_sojae_140303        ------------------------------------------------------------	253
PPTG_16278            IN----------------------------------------------------------	259
PPTG_19219            ------------------------------------------------------------	249
PITG_08944            ------------------------------------------------------------	240
PPTG_16271            ------------------------------------------------------------	240
P_capsici_15451       ------------------------------------------------------------	226
P_sojae_140289        ------------------------------------------------------------	199
P_sojae_109280        ------------------------------------------------------------	240
PPTG_16267            ------------------------------------------------------------	240
P_capsici_106898      ------------------------------------------------------------	239
P_sojae_140300        ------------------------------------------------------------	243
PITG_16991            ------------------------------------------------------------	243
PPTG_16273            ------------------------------------------------------------	242
P_capsici_107440      ------------------------------------------------------------	240
PITG_13560            ------------------------------------------------------------	205
P_sojae_109281        ------------------------------------------------------------	241
PPTG_16272            ------------------------------------------------------------	241
P_capsici_107334      ------------------------------------------------------------	241
PPTG_19220            ------------------------------------------------------------	271
P_capsici_15453       ------------------------------------------------------------	265
P_capsici_107122      ------------------------------------------------------------	265
P_capsici_15454       ------------------------------------------------------------	265
P_sojae_140302        ------------------------------------------------------------	251
PITG_16992            ------------------------------------------------------------	239
P_sojae_140301        ------------------------------------------------------------	246
P_capsici_107461      ------------------------------------------------------------	240
PITG_14060            ------------------------------------------------------------	226


PEBEL_30235           ------------------------------------------------------------	274
PEBEL_30158           ------------------------------------------------------------	267
HpaG808599            ------------------------------------------------------------	251
HpaG814377            ------------------------------------------------------------	244
P_capsici_7335        ------------------------------------------------------------	236
HpaG805382            --ATK-ASP--------------------------------------D-DETAEPGADEK	314
P_sojae_138787        GSSEE-QTPSTPSTGSSTEQTTDAPA--------------ASN-TGSS-EETPSTGSSEE	345
P_capsici_99469       GSSTQ-ETPSTSSTGSS------------------EQTPATSSTGSST-EETPSSGSSEE	347
PPTG_16566            GSSTE-QSPSTPSTGSSTEETPSASSTGSS----TEQNPSTPSTGSST-EETPSTGSSTE	361
PITG_06962            ---TE-E---TPSTGSSTEETPSTGSSTEETPSTGSSTENTPSTGSST-EETPTTGSSAE	359
PPTG_19378            ------------------------------------------------------------	238
P_sojae_144813        ------------------------------------------------------------	202
P_capsici_111741      ------------------------------------------------------------	241
P_sojae_109681        ------------------------------------------------------------	241
PPTG_19377            ------------------------------------------------------------	241
PPTG_16567            YSQGEMAASAAS--GTASS-------------VSGEAATN--NTTS-SVNAAASTSSSQS	320
P_sojae_119627        YSQQETTAPS----TS---------------------------------------STSVS	311
P_capsici_98553       YSTEETTTSSTPTTTTATS-------------VDGEATTS--IASSLSGETMTSTTSSQN	333
P_sojae_140303        ------------------------------------------------------------	253
PPTG_16278            ------------------------------------------------------------	259
PPTG_19219            ------------------------------------------------------------	249
PITG_08944            ------------------------------------------------------------	240
PPTG_16271            ------------------------------------------------------------	240
P_capsici_15451       ------------------------------------------------------------	226
P_sojae_140289        ------------------------------------------------------------	199
P_sojae_109280        ------------------------------------------------------------	240
PPTG_16267            ------------------------------------------------------------	240
P_capsici_106898      ------------------------------------------------------------	239
P_sojae_140300        ------------------------------------------------------------	243
PITG_16991            ------------------------------------------------------------	243
PPTG_16273            ------------------------------------------------------------	242
P_capsici_107440      ------------------------------------------------------------	240
PITG_13560            ------------------------------------------------------------	205
P_sojae_109281        ------------------------------------------------------------	241
PPTG_16272            ------------------------------------------------------------	241
P_capsici_107334      ------------------------------------------------------------	241
PPTG_19220            ------------------------------------------------------------	271
P_capsici_15453       ------------------------------------------------------------	265
P_capsici_107122      ------------------------------------------------------------	265
P_capsici_15454       ------------------------------------------------------------	265
P_sojae_140302        ------------------------------------------------------------	251
PITG_16992            ------------------------------------------------------------	239
P_sojae_140301        ------------------------------------------------------------	246
P_capsici_107461      ------------------------------------------------------------	240
PITG_14060            ------------------------------------------------------------	226


PEBEL_30235           ------------------------------------------------------------	274
PEBEL_30158           ------------------------------------------------------------	267
HpaG808599            ------------------------------------------------------------	251
HpaG814377            ------------------------------------------------------------	244
P_capsici_7335        ------------------------------------------------------------	236
HpaG805382            EEEKEE-----------------------EE---------TADLEPESS-----------	331
P_sojae_138787        TPSTGSSEETPAT---------SDAGSS-EETPATSSTGSSEETPVESS-----TGSSEQ	390
P_capsici_99469       TPT-GSSNDTPS----------SDAGSSSEQTPSTSSTGSSAQTPPTPS-----TGSTGS	391
PPTG_16566            QTPSTASGDDDTTTSSGDETETSGAGDQTTTSGASEETTPSSTTPPTTSSSSEETPSTGS	421
PITG_06962            ETPSTGSYRADLSDFFRGR-----------------------------------------	378
PPTG_19378            ------------------------------------------------------------	238
P_sojae_144813        ------------------------------------------------------------	202
P_capsici_111741      ------------------------------------------------------------	241
P_sojae_109681        ------------------------------------------------------------	241
PPTG_19377            ------------------------------------------------------------	241
PPTG_16567            GETTASS-----TTSTWNEQG--SSGSSATSTT---------TTTSTASSTWTETTAPS-	363
P_sojae_119627        GEATTSS-----TSSTWNEQE--STAASASETTA--------PSTTSS--TSEETTSSS-	353
P_capsici_98553       DKSTTSS-----TSSTWNEQE--SNVTSTNSSGA--------TTTTMTSSSWAETASP--	376
P_sojae_140303        ------------------------------------------------------------	253
PPTG_16278            ------------------------------------------------------------	259
PPTG_19219            ------------------------------------------------------------	249
PITG_08944            ------------------------------------------------------------	240
PPTG_16271            ------------------------------------------------------------	240
P_capsici_15451       ------------------------------------------------------------	226
P_sojae_140289        ------------------------------------------------------------	199
P_sojae_109280        ------------------------------------------------------------	240
PPTG_16267            ------------------------------------------------------------	240
P_capsici_106898      ------------------------------------------------------------	239
P_sojae_140300        ------------------------------------------------------------	243
PITG_16991            ------------------------------------------------------------	243
PPTG_16273            ------------------------------------------------------------	242
P_capsici_107440      ------------------------------------------------------------	240
PITG_13560            ------------------------------------------------------------	205
P_sojae_109281        ------------------------------------------------------------	241
PPTG_16272            ------------------------------------------------------------	241
P_capsici_107334      ------------------------------------------------------------	241
PPTG_19220            ------------------------------------------------------------	271
P_capsici_15453       ------------------------------------------------------------	265
P_capsici_107122      ------------------------------------------------------------	265
P_capsici_15454       ------------------------------------------------------------	265
P_sojae_140302        ------------------------------------------------------------	251
PITG_16992            ------------------------------------------------------------	239
P_sojae_140301        ------------------------------------------------------------	246
P_capsici_107461      ------------------------------------------------------------	240
PITG_14060            ------------------------------------------------------------	226


PEBEL_30235           -----------------------------------------	274
PEBEL_30158           -----------------------------------------	267
HpaG808599            -----------------------------------------	251
HpaG814377            -----------------------------------------	244
P_capsici_7335        -----------------------------------------	236
HpaG805382            -AGKEE----P-DSTSADVTSATIAIPTVKSKCILRRN---	363
P_sojae_138787        NPGQQE-PSTPTTSSSSEETP----STETNPKCVLRRVRRE	426
P_capsici_99469       STDQQETTTAPTTSSSSEETP----STGTNPKCTLRRVRRD	428
PPTG_16566            STGQQE----PTESSSSEETPSAPSTPATNPKCTLRRVRRD	458
PITG_06962            -----------------------------------------	378
PPTG_19378            -----------------------------------------	238
P_sojae_144813        -----------------------------------------	202
P_capsici_111741      -----------------------------------------	241
P_sojae_109681        -----------------------------------------	241
PPTG_19377            -----------------------------------------	241
PPTG_16567            ----------ASISTVESTTTSPTTSTITGSKCASRRVRRA	394
P_sojae_119627        ----------ATETTTTSVTAAPSTSTTTGKKCASRSVRRV	384
P_capsici_98553       ----------STTGASTTTASSVGTSTTTGQKCVSRRVRRV	407
P_sojae_140303        -----------------------------------------	253
PPTG_16278            -----------------------------------------	259
PPTG_19219            -----------------------------------------	249
PITG_08944            -----------------------------------------	240
PPTG_16271            -----------------------------------------	240
P_capsici_15451       -----------------------------------------	226
P_sojae_140289        -----------------------------------------	199
P_sojae_109280        -----------------------------------------	240
PPTG_16267            -----------------------------------------	240
P_capsici_106898      -----------------------------------------	239
P_sojae_140300        -----------------------------------------	243
PITG_16991            -----------------------------------------	243
PPTG_16273            -----------------------------------------	242
P_capsici_107440      -----------------------------------------	240
PITG_13560            -----------------------------------------	205
P_sojae_109281        -----------------------------------------	241
PPTG_16272            -----------------------------------------	241
P_capsici_107334      -----------------------------------------	241
PPTG_19220            -----------------------------------------	271
P_capsici_15453       -----------------------------------------	265
P_capsici_107122      -----------------------------------------	265
P_capsici_15454       -----------------------------------------	265
P_sojae_140302        -----------------------------------------	251
PITG_16992            -----------------------------------------	239
P_sojae_140301        -----------------------------------------	246
P_capsici_107461      -----------------------------------------	240
PITG_14060            -----------------------------------------	226

Hpa Hyaloperonospora arabidopsidis
PEBEL Peronospora belbahrii
P_capsici Phytophthora capsici
PITG Phytophthora infestans
PPTG Phytophthora parasitica
P_sojae Phytophthora sojae
The two catalytic residues in Phytophthora sojae protein 109681 that are essential for xyloglucan-degrading endoglucanase activity were highlighted in red (from Ma et al. 2015. Plant Cell 27: 2057-2072).

PEBEL_30158 and PEBEL_30235 were identified by Thines et al. 2020. MPMI 33: 742-753
All other proteins were taken from Ma et al. 2015. Plant Cell 27: 2057-2072
```
