## Supplementary material for "Dual transcriptional analysis of *Peronospora belbahrii* and *Ocimum basilicum* in susceptible interactions": SUPP FILE11

Characteristics of SWEET and ST proteins of *P. belbahrii*.

| Protein | Amino acids |  | MW^a^ | pI ^a^ | Subcellular location^b^ |
| --- | --- | --- | --- | --- | --- |
| SWEET transporters |  |  |  |  |  |
| belSWEET1^c^ | 241 |  | 26497 | 6.7 | Cell membrane |
| belSWEET2^c^ | 275 |  | 30390 | 6.8 | Cell membrane |
| PEBEL_05833 | 254 |  | 27974 | 9.7 | Cell membrane |
| PEBEL_04308 | 273 |  | 29796 | 7.2 | Cell membrane |
| PEBEL_06042 | 333 |  | 36441 | 6.3 | Cell membrane |
| PEBEL_04309 | 277 |  | 30212 | 7.9 | Cell membrane |
| MFS transporters |  |  |  |  |  |
| PEBEL_06041 | 271 |  | 30542 | 9.6 | Cell membrane |
| PEBEL_02444 | 513 |  | 55905 | 6.3 | Cell membrane |
| PEBEL_01548 | 519 |  | 56921 | 8.1 | Cell membrane |
| PEBEL_02039 | 457 |  | 50709 | 6.2 | ER membrane |
| PEBEL_05539 | 493 |  | 53289 | 6.1 | Lysosome membrane |
| PEBEL_08234 | 579 |  | 62582 | 4.8 | Cell membrane |
| PEBEL_01210 | 530 |  | 57152 | 8.1 | Lysosome membrane |
| PEBEL_01634 | 519 |  | 56136 | 7.6 | Lysosome membrane |
| PEBEL_00785 | 493 |  | 53813 | 6.5 | Lysosome membrane |
| PEBEL_01201 | 556 |  | 60856 | 6.2 | Lysosome membrane |
| PEBEL_00624 | 469 |  | 52461 | 6.1 | ER membrane |
| PEBEL_04315 | 510 |  | 55676 | 8.9 | Lysosome membrane |
| PEBEL_03529 | 494 |  | 53931 | 5.9 | Lysosome membrane |
| PEBEL_07073 | 520 |  | 57202 | 7.0 | Lysosome membrane |
| PEBEL_04194 | 465 |  | 49905 | 4.5 | Lysosome membrane |
| PEBEL_01212 | 539 |  | 59128 | 7.5 | Lysosome membrane |

^a^ calculated by Prot Pi protein tool; ^b^ predicted by DeepLoc-1.0; ^c^ could not be mapped to the *P. belbahrii* genome, but the putative CDS is on the next page

belSWEET1

ATGACAGATTTACTAACTGTAGCTATTGTACGTGTATTAGCCTCTTTAGCTGCATGCATCTTATTTGCAAGTCTATTACCGTCCATTAGGATTGTACACCAACGTAAAAGTACTGCAAGTATGCCCAGTGTACTTCCTGTATTGTCCATGATAGCAAATTGCATCACGTGGGGACTCTATGGCGTACTCATTGACGACTTTTTTCCATTAGTATTTACTAATATCATTGGAATATTCCTCTCACTCTTTTATCTCATTGTGTACTACTACTATGCGACGAACAAGGGTAGTTTAAGTCTTGAGATTCTAGCAACAGCTCTAGTGCTTGTGGGTATTTTACTATATCCATTGATGGCAGCTTTTGACAGGGTACAAGATGAAACCGTAGAGCACGTTGTGGGATTTCTTGCAGTAACTTCTTCGGCCATTATGTTTGGCTCGCCGCTGGTTCTAGTCAAAAGGGTGATTCGAGAACGCAACGCGAGTTCATTACCATTGAATATGATTACCACTGGAGTAGTCAATTGTGTCTTATGGCTTTGGTACGGTCTCTTGCTTGAAGACGCGTTCGTCATTGTACCGAATGCGGCAAATTTGCTTCTTGGCACCGTACAACTTGGACTCTTTTGCATTTATCCACGTAGCGGAACGTGCAACAACGTTGATTCTTCTACTTCTAAATTGAAACCCGAAGATAAAAAAGTACGGCAGATAGAAACGGAGTGA

belSWEET2

ATGGTCAACCTTATTGTAGAAACGCTCTTTCGTGTTTTAACGTCCGTTGCGTCTGTCAGTGTAACGTTGAGCATGGGTCCGTCAATCTACCGGATCTATTGCAACAGGGATACAGGTGTTGCGTCCGTGCTACCACTTGTATGTATGGTTACAAACGCCCATGTGTGGATGTTAAATGGTGTTACGGTTAAGAACTGGTTTCCCGTGTTTACCACCTTTATCACGTCTGACGTTATCGCTATATGTTACGTGACTGTATTCTACTGCTATGCGCGTGATCGTAAGAAAGCTCTTTGCAGGATCATTATTGGAAGTATTATTCTCTGTCTCGTCACGACCTACGCAATCCTGGGCAACGCAGGGCGCACCAATCAATCGCAAGATGGCGTAGAAACGACATTGGGTATCTTGGGTGTTTTGGCTGGGCTTAGCATGTTTTGCTCGCCATTCGAACGAATGATGAAAATACTACACTACAAGTCGGCGGTCTTTATCCCAATCCCGATGGTGGTGACTGGCACATTGAACAACCTTATGTGGCTCGTATACTGTCCAATGGTTGGAAGCTGGTTTCTCTTCGGAAGTAATGTAATGTGTCTGCTCGTGTGCTCGGTGAACCTTATTCTCTACATCATTTACAACCCAGAGACGCATCCGTTGCCCTTGGAAGAAGGTGATGGAGACGAAAACACTCCTTCTAGAGTCGAGTCAGTGTCCATCAATAAGGTACTTTCGCCGCAAGATACCGCTTTGAAAACGAAGCCAAACTTGTTCAGTCCAGCGTACGATCTACATCAGTCACCAGTTGCAGGCAATTTGACCGAATGA
